# Directionality of transcriptional regulatory elements

**DOI:** 10.1101/2024.12.01.621925

**Authors:** You Chen, Sagar R. Shah, Alden K. Leung, Mauricio I. Paramo, Kelly Cochran, Anshul Kundaje, Andrew G. Clark, John T. Lis, Haiyuan Yu

## Abstract

Divergent transcription is a critical marker of active transcriptional regulatory elements (TREs), including enhancers and promoters, in mammals. However, distal elements with unidirectional transcriptional patterns are often overlooked, leaving their identity and function poorly understood. Here, we performed a systematic comparison between divergent and unidirectional elements, revealing their distinct architectural and functional features. Our analysis also shows that unidirectional elements have younger sequence ages and are under weaker evolutionary constraints than divergent elements, indicating that they may represent a unique category of genomic regulatory function with more recent origins. Notably, we observed that some transcription factors, including CTCF, AP1, SP, and NFY, exhibit dual roles in modulating the directionality of TREs, either activating or repressing nascent transcription in a position-dependent manner. Overall, the elucidation of directionality enhances our understanding of the diverse architectural models, functional features, evolutionary dynamics, and regulatory logic of TREs.

## Introduction

Promoters and enhancers are two essential classes of transcriptional regulatory elements (TREs) that play crucial roles in modulating the genome’s regulatory programs in normal and disease states. Both elements can generate extensive transcriptional activity, and understanding their molecular architecture is crucial to uncovering their biological roles.

Active mammalian promoters often exhibit divergent transcription, resulting in both a stable sense transcript that represents the annotated gene, and an upstream antisense transcript that cannot elongate efficiently beyond the promoter region^1,2^. Despite their instability, low expression level, and lack of coding potential, a recent study has suggested that in progesterone gene regulation, the antisense transcripts may act as gatekeepers of promoter-proximal pause release, rather than being passive by-products of transcription^3^.

The directionality of promoters can differ across various human tissues and conditions. A study of 376 samples from nine different tissue types, including cancer and benign conditions, revealed lineage-specific and cancer-specific antisense transcription^4^. Moreover, well-positioned CTCF and cohesin have been demonstrated as a barrier against antisense transcription initiation at divergent promoters^5,6^. An explainable machine-learning model of transcription initiation, Puffin^7^, also identified only a few motifs with orientation-specific effects at promoters, including TATA-box and YY1.

In contrast to promoters, enhancers produce short-lived RNAs called enhancer RNAs (eRNAs)^8^. The presence of divergent transcription has proven to be a more robust predictor of active enhancers than classical epigenomic signatures (e.g., high chromatin accessibility, enrichment of H3K4me1 and H3K27ac)^9,10^.

A study of eRNA transcription in human lymphoblastoid cell lines derived from the Yoruba population identified approximately 10,000 genetic variants associated with the directionality of divergent enhancers, termed directional initiation quantitative trait loci (diQTLs)^11^. These diQTLs are significantly enriched in the vicinity of enhancer transcription start sites (TSSs), potentially affecting the core promoter motifs such as Initiator (Inr) and TATA-like elements. Moreover, the study showed that enhancer diQTLs are significantly associated with changes in gene expression.

While divergent transcription is a well-documented phenomenon in bulk transcriptome data, a recent study using C1 CAGE profiling has challenged this paradigm by suggesting that eRNA transcripts may arise from either strand in a mutually exclusive manner within individual cells^12^. However, the low sequencing coverage limited the detection of eRNAs, as exemplified by the fact that this study identified only 32 divergent enhancer loci with at least 10 reads in a minimum of 5 cells. Despite these technical limitations, the occurrence of unidirectional transcription at enhancers remains an intriguing phenomenon, warranting deeper investigation to fully understand its underlying mechanisms and biological significance.

Recently, a comprehensive evaluation of RNA sequencing approaches demonstrated that nuclear run-on with cap-selection (PRO-cap^13^) achieves the highest sensitivity and specificity for detecting eRNAs^14^. In this study, we leveraged the PRO-cap assay to identify a substantial number of unidirectional elements with sufficient sequencing depth. This enabled a systematic comparison of divergent and unidirectional elements in terms of their architectural features, functional characteristics, and evolutionary dynamics. Furthermore, we explored genetic contributors of transcriptional directionality shared by both enhancers and promoters, including CTCF and other directional motifs.

## Results

### Unidirectional elements have distinct architectural features

To examine transcriptional directionality at human enhancers and promoters, we performed PRO-cap, which captures RNA polymerase II initiation sites during nascent transcription at base-pair resolution, on two replicates of the chronic myelogenous leukemia cell line K562. Using our recently developed tool, Peak Identifier for Nascent Transcript Starts (PINTS)^14^, we identified a comprehensive set of active TREs genome-wide and categorized them into divergent elements (*n=*34,031) having a pair of peaks on opposite strands within 300 bp of each other, and unidirectional elements (*n*=19,346) with peaks on only one strand (Fig. 1a,b). To ensure a fair comparison between divergent and unidirectional categories, we applied a stringent set of criteria (see Methods for details) to derive elements for the analysis presented in this manuscript (Extended Data Fig. 1a,b).

**Fig. 1.**
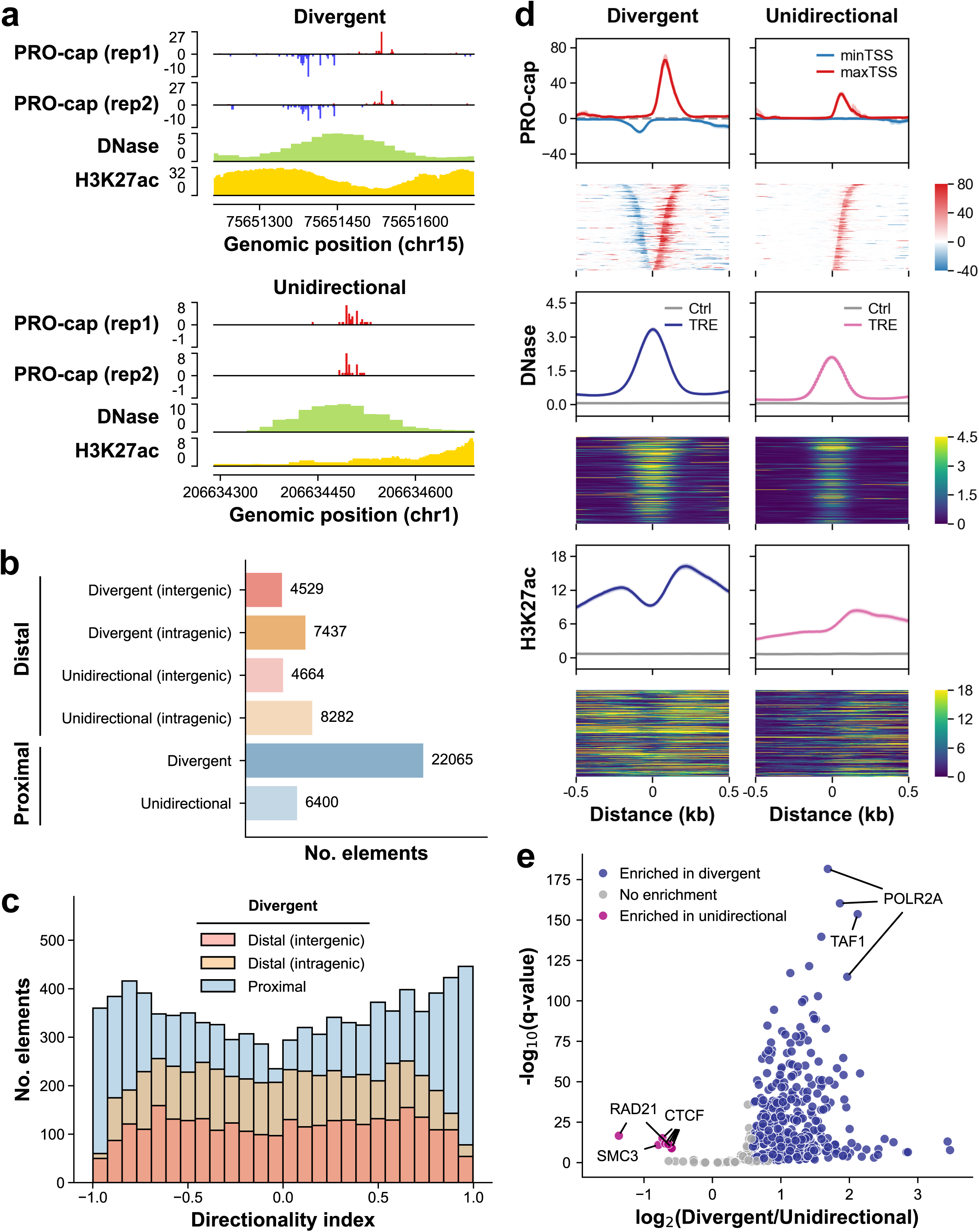
| Architectural features of divergent and unidirectional distal elements. **(a)** Genome browser tracks of PRO-cap, DNase-seq, and H3K27ac ChIP-seq for a representative divergent (top) and unidirectional (bottom) distal element. PRO-cap tracks from two replicates are shown, with red and blue signals indicating forward and reverse strands, respectively. **(b)** Genome-wide distribution of divergent and unidirectional elements identified by PRO-cap across proximal and distal intra- and intergenic regions. **(c)** Distribution of directionality index for divergent elements located in proximal and distal intra- and intergenic regions. **(d)** Metaplots and heatmaps of PRO-cap, DNase-seq, and H3K27ac ChIP-seq signals at divergent and unidirectional distal elements. The metaplots display average signals with 95% confidence intervals for each assay, while the heatmaps show signals at individual elements, sorted by distance from the prominent TSS to the center. **(e)** Volcano plot displaying ChIP-seq signal enrichment of DNA-binding proteins between divergent and unidirectional distal elements.

Beyond a binary classification of transcription patterns at TREs, we assessed the transcriptional directionality of divergent elements using a continuous metric, the directionality index (DI). This index is derived from the difference between PRO-cap read counts on the plus and minus strands relative to genomic coordinates, normalized by the total read counts for each element, revealing a spectrum of directionality. While some divergent elements exhibit similar transcription levels on both strands, others show pronounced, unbalanced transcription on one strand, skewing their transcriptional directionality (Fig. 1c). Notably, proximal elements tend to display higher DI compared to distal elements.

To explore the architectural features of these divergent elements, we positioned the TREs at the midpoint between two prominent TSSs on the plus and minus strands (Extended Data Fig. 1a). Both distal and proximal divergent elements were centered on DNase peaks and marked by the histone modification H3K27ac at their boundaries (Fig. 1d, Extended Data Fig. 1c). In the absence of distinct peaks detected on the opposite strand, unidirectional elements were anchored at the center of overlapping DNase peaks as a proxy (Extended Data Fig. 1a). We observed that transcription initiation at divergent and unidirectional elements occurs at variable distances from the center of open chromatin regions, with histone modification marks predominantly enriched on the same side that exhibits higher PRO-cap signals. Moreover, unidirectional elements have significantly lower signals for nascent transcription (PRO-cap) and epigenomic features (DNase-seq and H3K27ac ChIP-seq) compared to their divergent counterparts (Fig. 1d, Extended Data Fig. 1c).

To investigate protein binding patterns that may distinguish unidirectional from divergent elements, we analyzed ChIP-seq data from 311 target proteins and observed significant differences (Fig. 1e, Extended Data Fig. 1d). Divergent elements show greater enrichment for most proteins surveyed, including RNA polymerase II and general transcription factor components, consistent with their elevated transcription levels measured by the PRO-cap assay. In contrast, CTCF and cohesin components (RAD21 and SMC3) exhibit stronger binding signals in unidirectional elements, suggesting their potential involvement in establishing transcriptional directionality. Beyond differences in binding strength, some proteins (e.g., PHF8 and SMAD5) are bound on both sides of divergent elements, while they primarily bind to the side where PRO-cap peaks are detected in unidirectional elements (Extended Data Fig. 1e). Collectively, these results demonstrate that unidirectional elements have architectures distinct from divergent counterparts.

Beyond the overall architecture, we zoomed into the sequence content of individual transcriptional units within each category. Examination of single-nucleotide sequence preferences reveals two symmetric arcs of enriched AT content in divergent elements, whereas a similar pattern was detected on one side of unidirectional elements where PRO-cap peaks were observed (Fig. 2a). Closer inspection shows that each arc has a fainter parallel inner arc. Position-specific 3-mer frequencies indicated that the inner and outer arcs represent TATA-box-like and Inr-like sequences, respectively (Fig. 2b). Direct motif scans of TATA-box, Inr, and downstream promoter region (DPR) relative to prominent TSSs revealed that unidirectional TSSs and maximum TSSs (prominent TSS on the side with more read counts) of divergent elements have a slightly higher proportion of these motifs compared to minimum TSSs (prominent TSS on the side with less read counts) of divergent elements (Fig. 2c). Therefore, despite differences in their overall architectures, divergent and unidirectional elements have similar sequence content in individual core promoter regions.

**Fig. 2.**
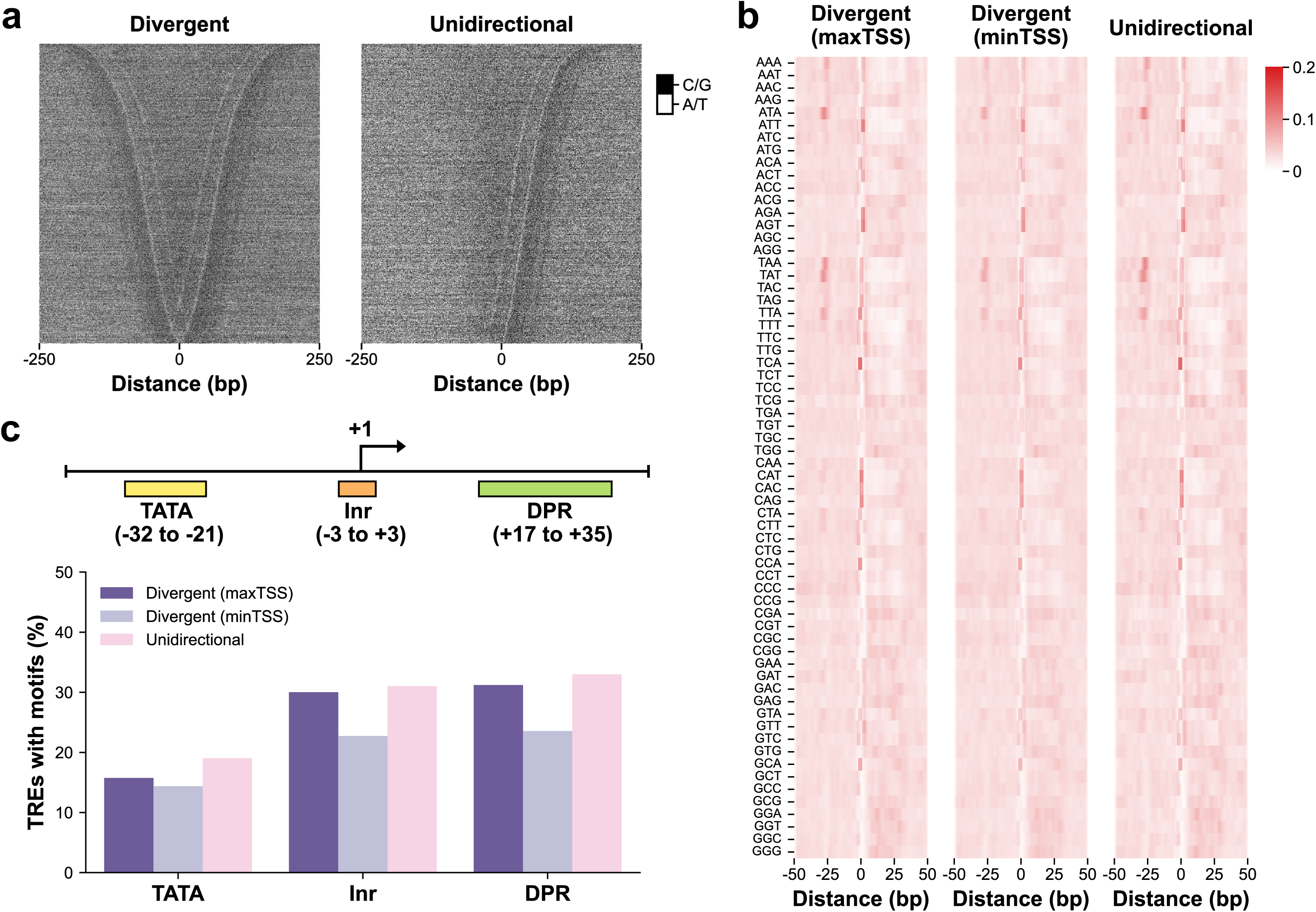
| Core promoter sequences at divergent and unidirectional distal elements. **(a)** Genomic DNA sequence content of divergent and unidirectional distal elements, with bases “A” and “T” colored in white, and bases “C” and “G” colored in black. The heatmaps show signals at individual elements, sorted by distance from the prominent TSS to the center. **(b)** The frequency of 64 three-mers at each position relative to the TSS, normalized to their total frequency within the region spanning −50 to +50 bp around prominent TSSs of divergent and unidirectional distal elements. **(c)** Percentage of elements displaying core promoter motifs matching the TATA box, Inr, or DPR in divergent and unidirectional distal categories.

### Potential confounders for binary classification

While our analysis thus far supports the discovery of unidirectional elements as a distinct class of TREs, we sought to rule out potential confounders that may impact this binary classification. First, we examined whether the absence of peaks on the other side of unidirectional elements resulted from poor sequence mappability. To that end, we evaluated the mappability using k-mers of varying lengths (24, 36, and 50 bp)^15^ based on the range of read lengths detected (Extended Data Fig. 2a). Both divergent and unidirectional elements have better mappability compared to the random genomic background. Importantly, no apparent bias was observed in unidirectional elements on the side lacking PRO-cap peaks (Extended Data Fig. 2b).

Insufficient sequencing depth could also potentially account for the missing peak at unidirectional elements. To address this possibility, we downsampled our PRO-cap dataset of ∼44 million (M) uniquely mapped PCR-free reads to lower read counts (30, 20, 10, 5, and 1 M) and performed peak calling on each downsampled dataset. We found that at extremely low depth (<10 M reads), a significant number of divergent elements are misclassified as unidirectional elements. These misclassified elements exhibit substantial transcription detected on the opposite strand at the original, higher sequencing depth (Extended Data Fig. 3a,c), indicating that a sufficient sequencing depth is critical for accurate discrimination of unidirectional from divergent elements. Therefore, to further explore the optimal sequencing depth required, we utilized a deep learning model, ProCapNet^16^, to assess transcription potential based on DNA sequences. The model was trained using peaks called at the original sequencing depth, with the exclusion of those overlapping with unidirectional elements identified at downsampled levels (profile task correlation: 0.520±0.006; count task correlation: 0.774±0.017). The results of the predicted PRO-cap signals from unidirectional elements at various read depths show that a minimum sequencing depth of ∼20 M reads is necessary to achieve an accurate binary classification (i.e., divergent versus unidirectional elements) of transcription directionality (Extended Data Fig. 3b,d). Taken together, our findings demonstrate that unidirectional elements are a distinct set of regulatory elements with unique transcriptional and architectural patterns and these TREs are not solely a product of experimental artifacts, poor mappability, or insufficient sequencing depth.

### A potential evolving process underlying directionality

Given the differences in overall architecture (i.e., one versus two transcription units) and the similarity in sequence content of individual transcription units, we hypothesized an evolving process from unidirectional to divergent distal elements. We found that, while both divergent and unidirectional distal elements are significantly older than random genomic background, unidirectional elements are slightly younger than divergent counterparts (Fig. 3a). In addition, in both TRE groups, the side with higher PRO-cap reads tends to be older and less tolerant to variation, as measured by context-dependent tolerance scores (CDTS)^17^. Furthermore, the phyloP^18^ patterns delineated open chromatin regions, showing the highest conservation at the center, with unidirectional elements exhibiting a narrower peak. Together, these results show that unidirectional elements have relatively younger sequence ages and weaker evolutionary constraints compared to their divergent counterparts, suggesting they represent a distinct class of functional genomic units with more recent evolutionary origins.

**Fig. 3.**
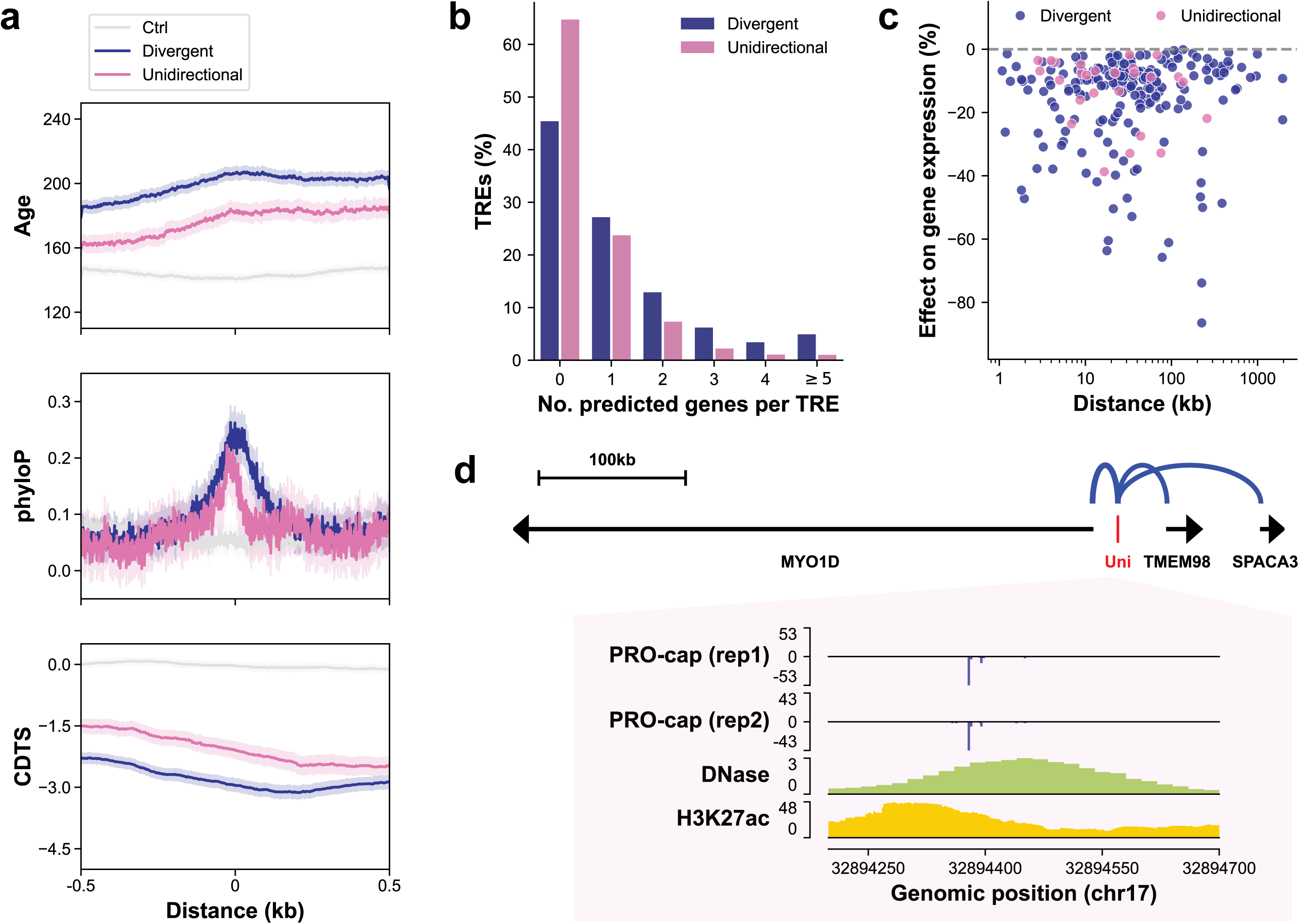
| Functional features of divergent and unidirectional distal elements. **(a)** Metaplots of sequence age (million years ago) and evolutionary metrics (phyloP and CDTS) across divergent and unidirectional distal elements. Distances are represented as ±0.5 kb from the center. **(b)** Histogram showing the number of genes predicted to be regulated by each divergent or unidirectional distal element as determined by the ABC model. **(c)** Percent change in target gene expression following CRISPR perturbations to divergent and unidirectional distal elements. **(d)** Schematic (top) illustrating the regulatory effects of a unidirectional distal element on three target genes (*MYO1D*, *TMEM98*, and *SPACA3*) validated by CRISPR experiments. Below are PRO-cap, DNase-seq, and H3K27ac ChIP-seq signal tracks for the unidirectional distal element.

Transposable elements (TEs) are well-documented as a prolific source of regulatory elements, playing a crucial role in shaping the human regulatory landscape^19,20^. To further explore the evolution of TREs, we first identified TE-derived distal elements by requiring that overlapping TE sequences be at least half the size of the TREs, as a conservative estimate. Approximately 25% of unidirectional elements, slightly more than divergent ones, overlap with TEs (Extended Data Fig. 4a). The proportion and enrichment of TE classes are similar between unidirectional and divergent elements (Extended Data Fig. 4b,c). Notably, long terminal repeat (LTR) elements are significantly enriched relative to their genomic proportion, likely because they often retain their original cis-regulatory sequences post-insertion in the genome^19^.

To further understand the potential evolutionary process from TEs to TREs, we examined their spatial relationships. If TEs randomly transition into TREs regardless of their insertion location, we would expect similar distance distributions between TEs overlapping with TREs and those not overlapping with TREs. Interestingly, we observed that TEs overlapping with TREs (d_2_ and d_3_) are significantly closer to non-TE divergent elements compared to TEs not overlapping with any TREs (d_1_), supporting the notion that genomic context can influence the likelihood that a TE acquires regulatory function (Extended Data Fig. 4d). TEs overlapping with unidirectional elements (d_2_), particularly those from LTR and long interspersed nuclear element (LINE) groups, are farther from non-TE divergent elements than TEs overlapping with divergent ones (d_3_). Together, our observations hint at a possible evolutionary process influencing directionality, where TEs may transition into unidirectional TREs and, over time, develop into divergent TREs.

### Unidirectional distal elements have gene regulatory functions

Considering their recent evolutionary origins, we aimed to determine whether unidirectional elements have acquired regulatory functions. We first examined their genomic distribution relative to genes and observed that unidirectional distal elements have similar genomic distributions to their divergent counterparts (Extended Data Fig. 5), implying that they may play a role in gene regulation.

To evaluate the regulatory potential of unidirectional distal elements, we employed the activity-by-contact (ABC) model^21^ to predict target genes in the K562 cell line. While a significant number of unidirectional distal elements is predicted to regulate target genes, a higher proportion of these elements have no predicted target genes than divergent distal elements (Fig. 3b). This result can be attributed to the lower activity of unidirectional distal elements, as indicated by DNase and H3K27ac ChIP-seq signals, despite having relatively comparable average 3D genome contact frequencies across all tested promoter-enhancer pairs with those involving divergent distal elements (Extended Data Fig. 6a). We further compared predicted target genes unique to either group or shared between both groups (Extended Data Fig. 6b). The unique and shared target gene sets are under similar selection strength as measured by loss-of-function observed/expected upper bound fraction (LOEUF) scores^22^, and exhibit comparable gene expression levels based on RNA-seq data (Extended Data Fig. 6c,d). Bergman *et al*^23^ identified two classes of promoters where P2 promoters showed stronger intrinsic activity and less responsiveness to enhancers. Among the target genes unique to unidirectional distal elements, there is a higher proportion of elements associated with P2 promoters, followed by those unique to divergent distal elements and shared ones (Extended Data Fig. 6e). Additionally, the promoters of these target genes unique to unidirectional distal elements display higher DNase and H3K27ac signals compared to the other two groups (Extended Data Fig. 6f).

In addition to *in silico* target gene predictions, we evaluated the regulatory effects of unidirectional distal elements using previously curated CRISPR perturbation experiments^24^. Although a larger fraction of divergent distal elements than unidirectional elements showed a significant ability to activate expression of cognate genes, 23 unidirectional distal elements were found to activate the expression of their cognate target genes (Fig. 3c). For instance, the deletion of one unidirectional element significantly reduced the expression levels of three target genes (*MYO1D*, *TMEM98*, and *SPACA*3) by 30-40% (Fig. 3d). Altogether, these results demonstrate that at least a subset of unidirectional distal elements possess gene regulatory function.

### CTCF binding blocks transcription initiation at TREs

Beyond the binary classification of divergent and unidirectional elements, we further investigated contributing factors underlying transcriptional directionality along a continuous spectrum. Noting our observation that CTCF and cohesin components are enriched in unidirectional elements in the K562 cell line (Fig. 1e, Extended Data Fig. 1d), we first examined whether these factors influence transcriptional directionality at TREs. A previous study by the Blobel lab demonstrated that CTCF regulates the directionality of divergent promoters by blocking antisense transcription^6^. To determine if CTCF also represses transcription initiation at enhancers, we performed PRO-cap in the human colorectal carcinoma cell line HCT116, both before and after inducing CTCF degradation using an auxin-inducible degron system.

In the unperturbed HCT116 cell line, we observed higher levels of CTCF and RAD21 ChIP-seq signals at unidirectional distal elements compared to divergent ones, consistent with our findings in the K562 cell line (Extended Data Fig. 7a). Further classifying divergent elements based on their absolute DI, our analysis demonstrated that highly skewed divergent elements exhibit stronger CTCF and RAD21 binding compared to more balanced ones (Extended Data Fig. 7b).

Next, we investigated the impact of acute CTCF depletion on transcription initiation at distal divergent and unidirectional elements. For subsequent analyses, the maximum TSS side is defined as the side with more read counts for divergent and unidirectional elements before CTCF degradation, while the opposite side is termed the minimum TSS side. We observed a significant increase in transcription levels at 223 distal elements on the minimum TSS side following CTCF loss (Fig. 4a, c), while the maximum TSS side remains largely unchanged (Extended Data Fig. 8a). The majority (∼95%) of these transcriptional perturbed elements are initially bound by CTCF, providing further evidence of CTCF’s direct role in transcriptional changes (Extended Data Fig. 7c). Notably, a higher proportion of unidirectional and transcriptionally skewed divergent elements, which have higher CTCF and RAD21 binding signals, show an increase in transcription after CTCF loss (Extended Data Fig. 7d). Furthermore, CTCF-bound elements in the upregulated group exhibit stronger CTCF and RAD21 binding (Extended Data Fig. 7e) and are enriched at chromatin loop anchors (Extended Data Fig. 7f) and chromatin domain boundaries (Extended Data Fig. 7g), compared to those in the unchanged group. Thus, our results indicate that CTCF occupancy at TREs blocks transcription initiation on one side, thereby tuning their transcription directionality.

**Fig. 4.**
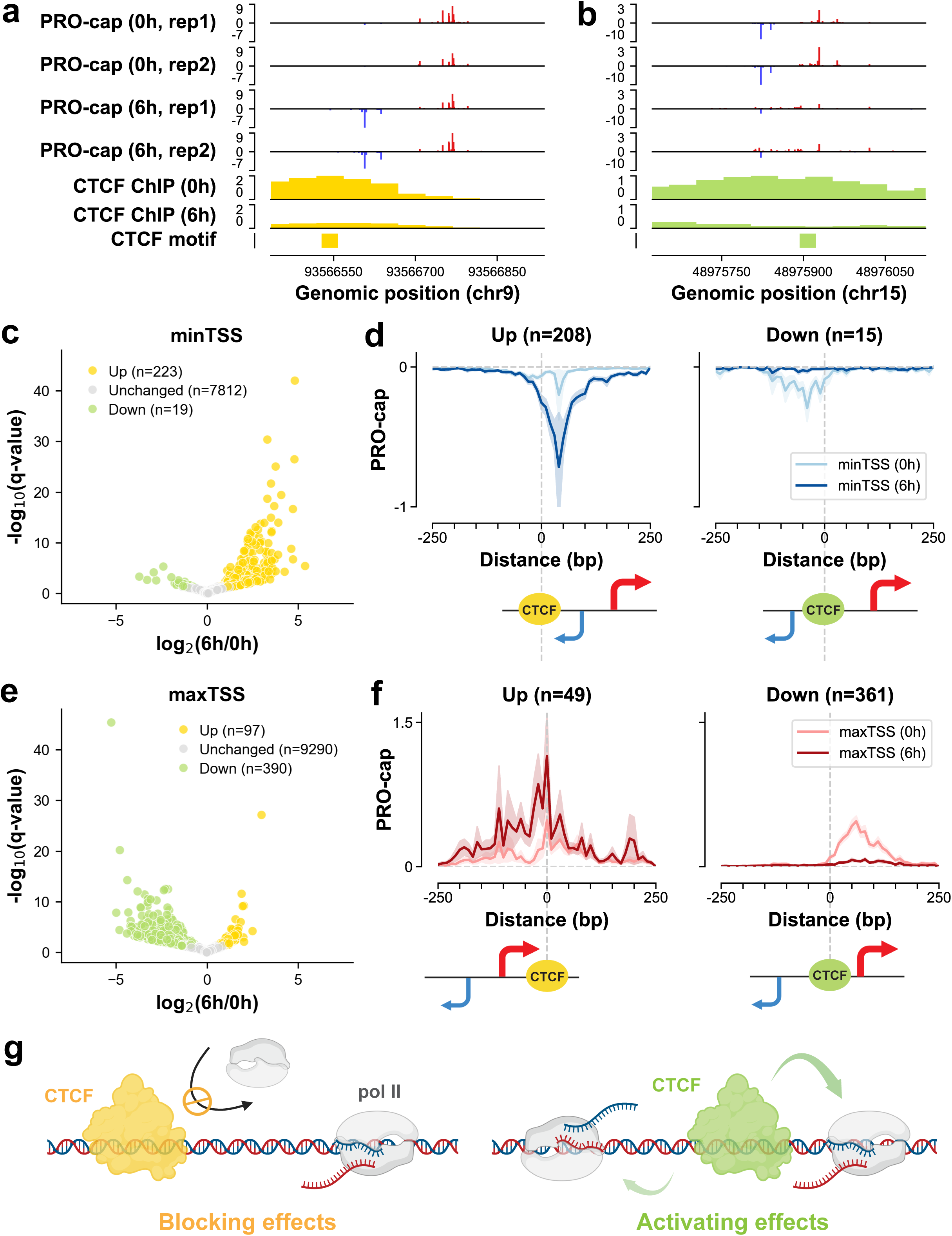
| CTCF has a dual function in modulating transcription initiation. **(a,b)** PRO-cap and ChIP-seq browser tracks of two representative distal elements, showing either upregulated (a) or downregulated (b) transcription before (0h) and after (6h) CTCF degradation. The positions of CTCF motifs are indicated and PRO-cap signal tracks from two replicates are shown. (c) Volcano plot of PRO-cap signals on the minimum TSS side of distal elements before and after CTCF degradation. Differentially expressed distal elements are highlighted in yellow for upregulation and green for downregulation in transcription. (d) Metaplots of PRO-cap signals (RPM normalized, 10-bp bins) on the minimum TSS side of distal elements in the upregulated (left) and downregulated (right) transcription groups, centered on CTCF motifs. A schematic of the CTCF motif position relative to transcription initiation sites is shown below each panel. (e) Same as (c), but for the maximum TSS side of distal elements. (f) Same as (d), but for the maximum TSS side of distal elements. (g) Illustration of CTCF’s dual function in modulating transcription initiation based on the motif’s position relative to transcription initiation sites.

### CTCF has a dual function in modulating transcription in a position-dependent manner

Alongside the changes in transcription levels, we noted an intriguing shift in TSS positions following the depletion of CTCF. For subsequent analyses, “downstream” denotes the direction in which transcription proceeds after a given TSS, while “upstream” refers to the opposite direction. In the upregulated group, some divergent elements exhibit an upstream repositioning of their minimum TSSs, moving closer to their respective maximum TSSs (Extended Data Fig. 9a). For instance, in one representative distal element, transcription initiates downstream of the CTCF motif before degradation. Upon CTCF loss, however, transcription initiates from positions upstream of the CTCF motif, suggesting that CTCF may normally block transcription of these cryptic TSSs (Extended Data Fig. 9b,c).

To further evaluate the relationship between CTCF function and its position relative to the TSS, we analyzed changes in transcription initiation focused on the minimum TSS side of TREs centered on the CTCF motif. We found that in the transcriptionally upregulated group, the CTCF motif is predominantly located downstream of the minimum TSS (Fig. 4d). Conversely, in the transcriptionally downregulated group, the CTCF motif resides in the upstream region, suggesting that CTCF has a dual function in tuning transcription initiation depending on its position relative to the original TSS. Similar trends were noted when analyzing changes in PRO-cap signals at the maximum TSS side (Fig. 4b,e,f). Moreover, our analysis of changes in the minimum and maximum TSS sides revealed a greater number of elements in the upregulated and downregulated groups, respectively (Fig. 4c,e). This pattern reflects a biological bias towards identifying CTCF as a blocker on the minimum TSS side, characterized by lower transcription levels, and as an activator on the maximum TSS side, characterized by higher transcription levels. Collectively, these findings indicate that CTCF functions as a transcriptional blocker when located downstream of the TSS and as an activator when positioned upstream (Fig. 4g).

Furthermore, we observed that elements with upregulated transcription at the minimum TSS side or downregulated transcription at the maximum TSS side show limited changes in transcription levels at their respective opposite sides (Extended Data Fig. 8a-d). Additionally, transcriptionally skewed elements become more balanced either by increasing transcription levels at the minimum TSS side or decreasing them at the maximum TSS side following CTCF depletion (Extended Data Fig. 8e,f). Similar patterns were observed in proximal elements (Extended Data Fig. 10), supporting a dual role for CTCF in regulating the transcription levels and directionality of both distal and proximal elements in a position-specific manner.

### Additional motifs critical for transcriptional directionality of TREs

Since CTCF can only account for a limited proportion of elements with skewed transcription patterns, we expanded our search for additional contributors of directionality in a more systematic manner. We trained the ProCapNet model using all elements found in the HCT116 cell line before CTCF degradation (profile task correlation: 0.600±0.004; count task correlation: 0.768±0.014) to identify motifs critical for transcription level (Extended Data Fig. 12a). We reasoned that motifs with orientation- and/or position-dependent effects on total PRO-cap read counts could be a source of unbalanced transcription between the two strands.

We first assessed whether the model has learned CTCF’s dual function in transcription initiation. We predicted PRO-cap signals before and after *in silico* deletion of CTCF motifs in elements with up- or down-regulated transcription levels in the CTCF degron experiments (using the same elements and motif positions as shown in Fig. 4d-left, f-right). The model demonstrated the ability to predict motif position-dependent changes consistent with our observations from the CTCF perturbation experiments, even though it was trained using only PRO-cap data obtained before CTCF degradation (Extended Data Fig. 11).

Next, we examined which motifs exhibited orientation-dependent effects on transcription levels. While most motifs do not show a clear orientation bias, some (TATA box and CTCF) are preferentially located on the same strand as the prominent TSSs of unidirectional elements (Extended Data Fig. 12b). The lack of bias observed for certain motifs, such as CREB and NRF1, can be attributed to their palindromic sequences. On the other hand, orientation preference may arise from structural constraints. For instance, the oriented binding of TATA box binding protein to the TATA box aligns with the assembly of a pre-initiation complex possessing a unique polarity^25^. Additionally, CTCF acts as a structural regulator of chromatin looping, where the orientation is crucial for loop topology and gene expression^26^.

We then tested whether other TFs might also exhibit a dual function in a position-dependent manner, similar to what was observed with CTCF. Since the ProCapNet model learned CTCF’s dual function solely from PRO-cap data before CTCF degradation, we reasoned that it might have also detected the dual function of other motifs, if any, without requiring PRO-cap data obtained after knocking down the corresponding TFs. We performed *in silico* deletion of motif instances and compared the fold change in transcription levels on either minimum or maximum TSS side (Fig. 5a). In addition to CTCF, we found that AP1, SP, and NFY also exhibit a dual function in modulating transcription at TREs (Fig. 5b). Specifically, these TFs are more likely to activate transcription when located upstream (decreased transcription after deletion) while repressing transcription when located downstream of TSSs (increased transcription after deletion). Two representative loci highlight the changes in predicted PRO-cap signals when deleting AP1 motifs downstream (Fig. 5c) or upstream (Fig. 5d) of the TSS, respectively. Taken together, motifs with orientation- and/or position-dependent effects on transcription are potential contributors to transcription directionality.

**Fig. 5.**
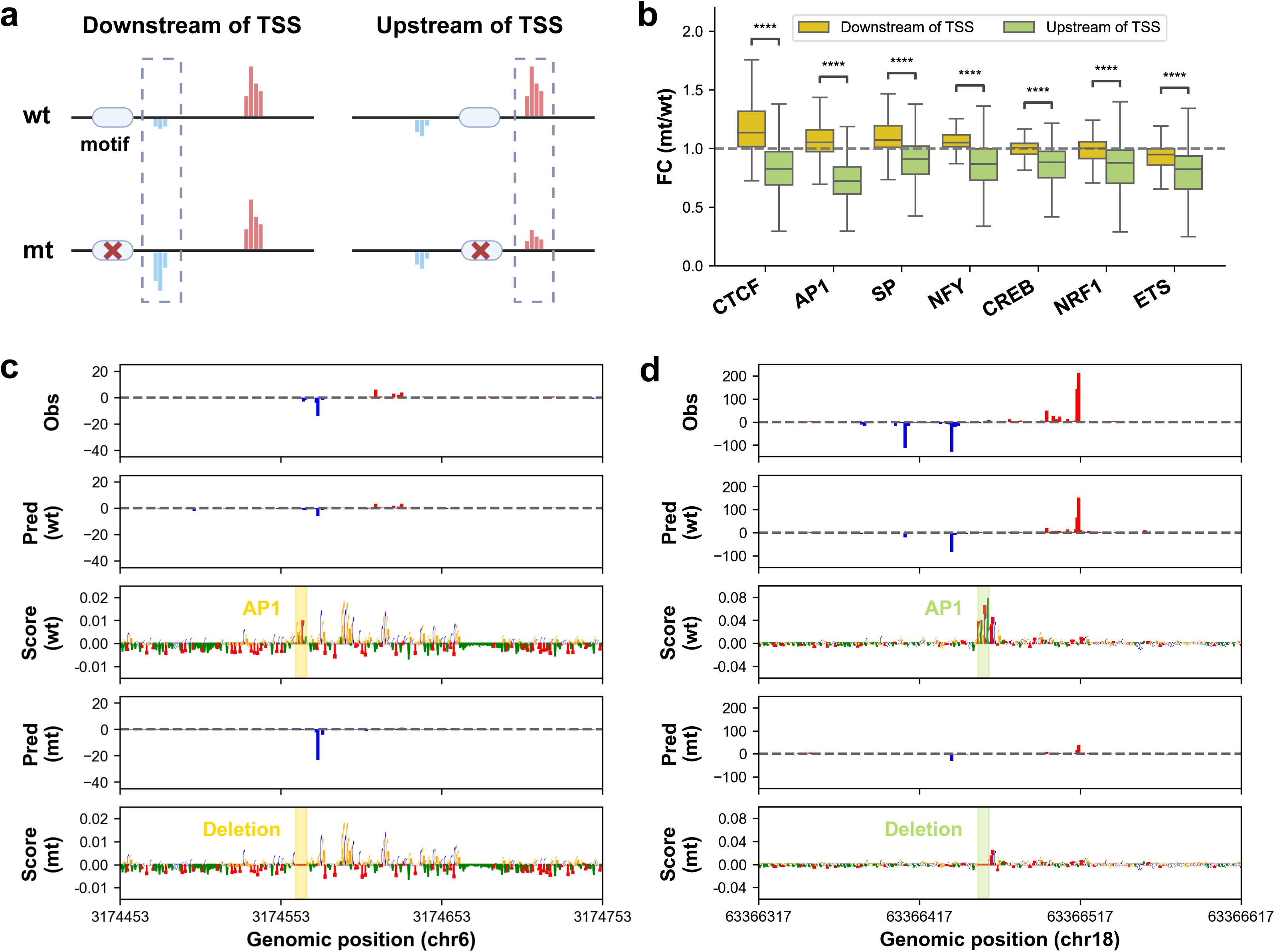
| TFs with dual functions in tuning transcription initiation. **(a)** A schematic of *in silico* motif deletion. **(b)** Barplots showing the fold change in predicted transcription levels at the minimum and maximum TSS sides following the *in silico* deletion of motif instances located downstream or upstream of the TSSs, respectively. **(c-d)** Representative loci where *in silico* deletion of AP1 motifs results in either transcription upregulation (c) or downregulation (d). The first track displays observed PRO-cap data, while the subsequent four tracks present predicted PRO-cap signals and their corresponding contribution scores from the count task based on original and mutant sequences. AP1 motifs are highlighted in shaded areas.

## Discussion

Our high-quality PRO-cap data, combined with our state-of-the-art peak calling tool PINTS, enabled us to identify a comprehensive list of divergent and unidirectional elements. Our systematic comparison of epigenetic features, protein binding profiles, and sequence content revealed that unidirectional elements are distinct biological entities with unique architectures compared to divergent elements. Furthermore, our data demonstrate that some unidirectional elements have acquired regulatory function, as shown by *in silico* target gene predictions and supported by CRISPR perturbation experiments. Our analyses also suggest a potential evolving process underlying the directionality of TREs. Future PRO-cap data from a diverse array of species may reveal further insights into this progression. Moreover, such data will enable the examination of how changes in directionality at TREs can affect the transcription levels of target genes, particularly those under positive selection that drive phenotypic adaptations over longer evolutionary timescales.

In addition to the evolutionary forces underlying directionality changes, we explored other contributors using the CTCF degron system coupled with PRO-cap, as well as ProCapNet deep learning models. We identified several TFs with dual functions in modulating nascent transcription based on their relative positions to the TSS. Interestingly, another group recently reported this spatial grammar through independent lines of functional evidence^27^, including TF perturbation (NRF1 and YY1 knockdown csRNA-seq data in human U2OS cells; NFY knockdown Start-seq data in mouse embryonic fibroblasts), natural genetic variation (csRNA-seq data in bone-marrow-derived macrophages from two mouse strains; PRO-cap dataset of 67 human lymphoblastoid cell lines), and TSS-MPRA. Our study adds to this growing body of evidence by characterizing a high-quality CTCF perturbation dataset with the following advantages^14^: 1) PRO-cap offers better sensitivity and specificity in eRNA detection compared to csRNA-seq and Start-seq; 2) PRO-cap can identify precise TSS positions of TREs; and 3) our study utilized PINTS, which outperforms HOMER, the peak-calling tool used in their study, in identifying active TREs genome-wide. Additionally, we employed deep learning models to decode the spatial grammar of other TFs, providing complementary evidence for position-dependent functional profiles.

Overall, our dissection of the mechanistic underpinnings of transcriptional directionality augment our understanding of the rules and logic governing gene regulation.

## Limitations

In an effort to systematically delineate the features of unidirectional TREs, we performed stringent filtering of the elements to ensure the quality and fairness of the comparative analyses. While this approach enhanced the clarity of our findings, it also led to the exclusion of certain elements that may warrant further investigation in the future. For example, unidirectional elements that do not overlap with DNase peaks were excluded from our analyses. However, this exclusion does not necessarily indicate that these elements are noise. Instead, it is plausible that their presence could reflect the higher sensitivity of the PRO-cap assay or differences in cell populations assayed by PRO-cap and DNase-seq experiments. Additionally, elements with other TREs within a 500 bp region were excluded from this study because we were unable to confidently dissect the contribution of individual TREs, which could have confounded our analyses. Future studies on these elements could yield new insights into the interaction and coordination among TREs, potentially uncovering complex regulatory networks. As a next step, to deepen our understanding of the relationship between transcriptional directionality and TRE activity, additional perturbations and functional experiments are essential to elucidate how these directional features influence gene regulation and cellular processes.

## Methods

### Preparation of PRO-cap libraries

Library preparations for two replicates, each with 10 million K562 cells, were conducted as previously described^13,28,29^. Likewise, we generated PRO-cap libraries from two replicates, each of 10 million HCT116 cells genetically modified using CRISPR targeting *H. sapiens* CTCF and *O. sativa* LOC4335696, before and after 6 h of treatment with 1 μM 5-phenyl-1H-indole-3-acetic acid. Briefly, the cells were permeabilized, followed by run-on reactions. After isolating RNA, two adaptor ligations and reverse transcription were performed using custom adaptors. 5’ cap selection reactions were conducted between adaptor ligations through a series of enzymatic steps. RNA washes, phenol:chloroform extractions, and ethanol precipitations were performed between reactions, all under RNase-free conditions. Libraries were sequenced on Illumina’s NovaSeq platform after PCR amplification and library clean-up. Details of library information can be found in Supplementary Table 1.

### Data preprocessing

This dataset was managed and analyzed using the Resource Management System (https://github.com/aldenleung/rmsp/). Raw reads were preprocessed with fastp^30^ (v0.23.4) for adapter trimming and unique molecular index (UMI) processing, retaining only reads ≥18 bp for downstream analyses. The processed reads were aligned to the human reference genome hg38 (GCA_000001405.15) and ribosomal DNA (U13369.1) using STAR^31^ (v2.7.11a). For the HCT116 PRO-cap library with Drosophila spike-in, the alignment also included the Drosophila reference genome (dm6). Uniquely mapped reads were filtered with samtools^32^ (v1.18) and deduplicated using umi_tools^33^ (v1.1.5). The PCR-free, uniquely mapped reads obtained from replicates of each sample were converted to bigwig format and merged with biodatatools (v0.0.7) for peak calling.

### Identification and classification of TREs

We performed peak calling with PINTS (v1.1.10), using the parameter “*--min-lengths-opposite-peaks 5*.” The identified peaks were then classified into two categories: (1) divergent peak pairs that consist of peaks located on opposite strands and within 300 bp of each other, and (2) unidirectional peaks that originated from only one strand. These elements were further divided into proximal (within ±500 bp) and distal (outside ±500 bp) categories based on their distances to TSSs of all genes annotated by GENCODE v37^34^.

All elements were resized to 500 bp to minimize bias toward large peaks in subsequent analyses. For divergent elements, the anchor point was defined as the midpoint between prominent TSSs on the plus and minus strands. For unidirectional elements, the center of overlapping DNase peaks was used as a proxy. To do so, the following inclusion criteria were specifically applied for unidirectional elements: (1) overlapping with DNase peaks; and (2) the center of the DNase peak must be appropriately positioned (i.e., on the left of the prominent TSS if the unidirectional element is on the forward strand, or on the right if on the minus strand). The resized divergent and unidirectional elements were further filtered using the following exclusion criteria: (1) elements extending beyond chromosome ends; (2) overlapping with ENCODE blacklist regions^35^; (3) original peak region not fully contained within the resized 500 bp region; and (4) overlapping with any other elements.

### Metaplots and heatmaps

To make the comparison and visualization more straightforward in the metaplots, the sides with more PRO-cap read counts were flipped to the right regardless of their relative orientation to genomic positions. The random genomic regions depicted in the metaplots were generated as controls with the following criteria: (1) matched on chromosome number and element size, and (2) not overlapping GENCODE v37 exon regions and ENCODE blacklist regions. In the heatmaps, elements were sorted based on the distance between their prominent TSSs and the anchor points. Any deviations from this sorting method were carefully noted.

### Annotating elements with protein binding features

To annotate elements with protein binding features, we downloaded a list of human TF ChIP-seq narrowpeak files in K562 cells from the ENCODE database^36^ and annotated each element with the maximum signalValue column for any overlapping peak (or 0 if there was no overlap). We then compared the fold-change (FC) of average signals between divergent and unidirectional categories. P-values were calculated using a two-sided t-test, followed by Benjamini-Hochberg correction. Enrichment was defined as |FC| > 1.5 and false discovery rate (FDR) < 0.05.

### Detection of core promoter elements

The presence of an initiator (Inr) was annotated for each prominent TSS when the sequence of −3 to +3 matched the consensus sequence BBCABW. The TATA box (−32 to −21 relative to the prominent TSS) and the downstream promoter region (DPR) elements (+17 to +35 relative to the prominent TSS) were identified using a previously published support vector regression model^37^.

### ProCapNet model training

We trained the ProCapNet models for each biosample using default parameters except that we set “in_window” as 1000, “out_window” as 500, and “source_fracs” as [1,0]. To ensure sufficient training dataset, we included all bidirectional and unidirectional elements (direct output from PINTS without additional filtering). For the model discussed in the section “Potential confounders for binary classification,” we additionally excluded elements that overlapped with unidirectional elements we aimed to predict at various downsampled levels. In the section “Additional motifs critical for transcriptional directionality of TREs,” we identified individual instances of the sequence patterns reported by TF-ModDISco^38^, following the procedure described by Cochran *et al*^16^. For follow-up analyses, we used the set of hits with high sequence- and contribution-match scores.

### Target gene prediction

We employed the ABC model to predict enhancer-gene interactions in the K562 cell line, following the protocol described in https://abc-enhancer-gene-prediction.readthedocs.io/en/latest/. A threshold of 0.025 was chosen as it achieved 70% recall in the CRISPR benchmark following the pipeline described in https://github.com/EngreitzLab/CRISPR_comparison. We assigned our PRO-cap distal elements to ABC regions that have an overlapping length of at least half the element size.

### Enhancer-gene interactions validated by CRISPR screens

To validate the regulatory effects of our distal elements, we intersected them with regions screened by CRISPR experiments curated at https://github.com/EngreitzLab/CRISPR_comparison, requiring an overlapping length of at least half the element size.

### Sequence age and evolutionary metrics

Although the age of sequences itself is not necessarily the age when the sequences first gained activity, it can serve as an upper bound. To estimate the sequence age, we employed the syntenic block aging strategy^39^, which calculates the time to the most recent common ancestor (MRCA) using a phylogenetic tree. We obtained the hg38 100-way vertebrate Multiz alignment data from the UCSC database^40^ and assigned an age to each syntenic block based on the MRCA of the species in the alignment. To simplify the analysis, we grouped MRCA nodes into 10 age categories and reported the age as the oldest ancestral branch. Given the high turnover of regulatory elements, phyloP and CDTS provide complementary information. phyloP scores were downloaded from the UCSC database to measure the conservation across species, while CDTS computed on 7,794 unrelated individuals was obtained to assess recent and ongoing purifying selection in the human lineage.

### Annotation of TE-derived TREs

Genomic TE annotations were obtained from RepeatMasker^41^. TE-derived distal elements were defined by requiring overlapping TE sequences to be at least half the size of the TREs, which serves as a conservative estimate. The enrichment of TE families for TREs was calculated as 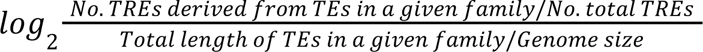.

### CTCF-related analysis

We only included elements present in the HCT116 cell line before CTCF degradation. All elements were resized to 500 bp, as described in “Identification and classification of TREs,” to quantify transcriptional changes in the maximum and minimum TSS sides. Differential expression analysis was performed using DESeq2^42^ with thresholds set as FDR < 0.05 and |FC| > 1.5 after shrinkage^43^. For elements with CTCF ChIP-seq binding, we used FIMO^44^ with default parameters to identify matches for the CTCF motifs (MA0139.2, MA1929.2, and MA1930.2) in the JASPAR database^45^. Only elements with confirmed CTCF ChIP-seq binding in HCT116 and high-confidence CTCF motifs were included to ensure precise positioning. We evaluated the proportion of CTCF motifs located at chromatin loop anchors using the CTCF chromatin interaction analysis by paired-end tag sequencing (ChIA-PET) data in HCT116 obtained from the ENCODE database. The insulation scores were calculated from HCT116 Hi-C data at a 10 kb resolution with a 30 kb window size using cooltools^46^. Locations with lower scores, indicating reduced contact frequencies between upstream and downstream loci, are often referred to as boundaries.

### Schematics

All schematics in Figs. 4, 5, and Extended Data Figs. 1, 4, and 9 were created using BioRender, with the appropriate publication licenses.

## Data availability

The PRO-cap datasets are deposited in ENCODE under accession numbers ENCSR165QKR (HCT116, CTCF_U), ENCSR261OWE (HCT116, CTCF_T), and ENCSR220XSM (K562). A summary of additional public datasets is provided at https://github.com/haiyuan-yu-lab/TRE_directionality/data_preprocessing/2.Other_resources.ipynb.

## Supporting information

all extended data figures

## Code availability

All code is available at https://github.com/haiyuan-yu-lab/TRE_directionality.

## Acknowledgements

We would like to thank all members of the ENCODE consortium for generating the datasets used in this paper and Jennifer Jou for assistance with data uploads to the ENCODE portal.

We acknowledge funding support from the National Institutes of Health (R01 DK115398 and R01 AG077899 to H.Y.; UM1 HG009393 and RM1 5RM1GM139738 to J.L. and H.Y.; 5U24HG007234, U01HG009431, U01HG012069, and U24HG007234 to A.K.). S.R.S. was supported in part by the Center for Vertebrate Genomics Scholar Award and the Breast Cancer Coalition’s Postdoctoral Fellowship. K.C. was supported in part by the Pierre & Christine Lamond Stanford Graduate Fellowship.

