## Supplementary material for "Directionality of transcriptional regulatory elements": all extended data figures

**Extended Data Figure 1 | Architecture of proximal elements**

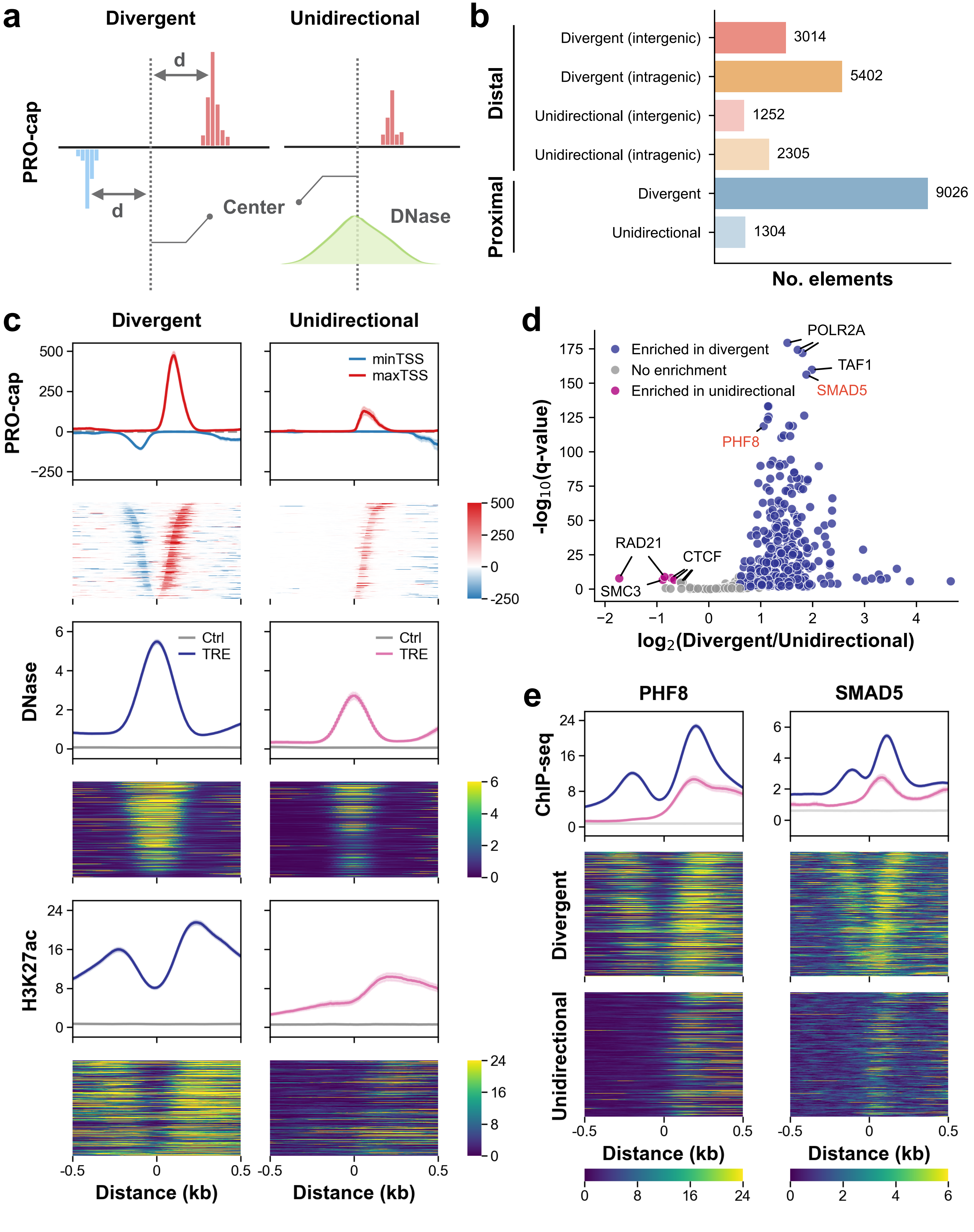

**Extended Data Fig. 1 | Architectural features of divergent and unidirectional proximal elements.**

**(a)** Diagram illustrating anchor points for divergent and unidirectional elements. For divergent elements, the midpoint between forward and reverse prominent TSSs serves as the anchor point, while for unidirectional elements, the center of overlapping DNase peaks is used.

**(b)** Genome-wide distribution of divergent and unidirectional elements identified by PRO-cap across gene proximal and distal intra- and intergenic regions after filtering steps (see Methods).

**(c)** Metaplots and heatmaps of PRO-cap, DNase-seq, and H3K27ac ChIP-seq signals at divergent and unidirectional proximal elements. The metaplots display average signals with 95% confidence intervals for each assay, while the heatmaps show signals at individual elements, sorted by distance from the prominent TSS to the center.

**(d)** Volcano plot displaying ChIP-seq signal enrichment of DNA-binding proteins between divergent and unidirectional proximal elements.

**(e)** Metaplots and heatmaps of PHF8 and SMAD5 ChIP-seq signals at divergent and unidirectional proximal elements. The metaplots display average signals with 95% confidence intervals for each assay, while the heatmaps show signals at individual elements, sorted by distance from the prominent TSS to the center.

### Extended Data Figure 2 | Sequence mappability

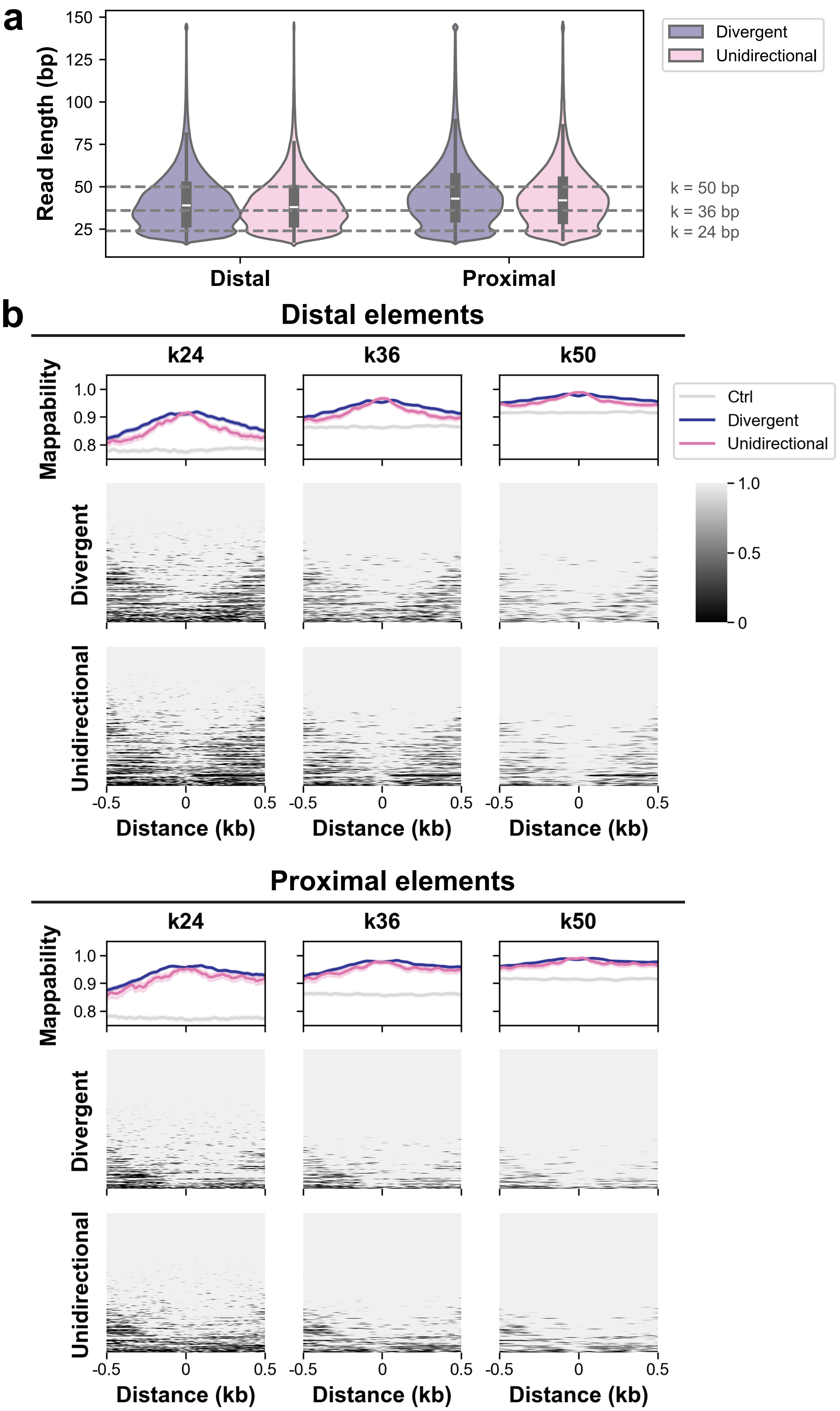

**Extended Data Fig. 2 | Sequence mappability at divergent and unidirectional elements.**
**(a)** Length distribution of reads mapped to divergent and unidirectional elements.
**(b)** Sequence mappability with k-mers of different lengths (24, 36, and 50 bp). The metaplots show the average mappability with 95% confidence intervals for each assay, while the heatmaps display the mappability at individual elements, sorted by the sum of 24-mer mappability. Distances are represented as ±0.5 kb from the center.

#### Extended Data Figure 3 | The effects of sequencing depth on binary classification

##### Distal elements

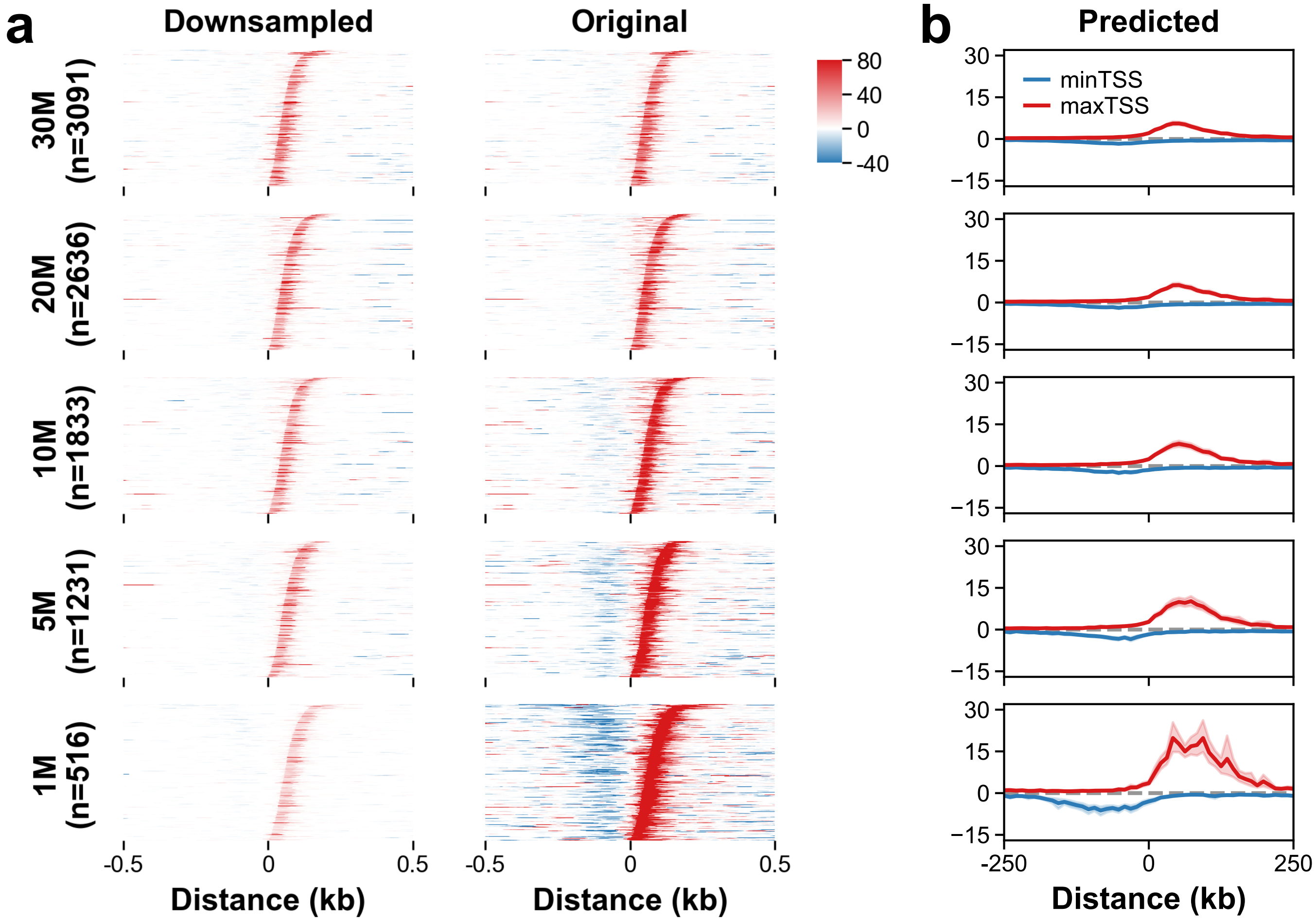


##### Proximal elements

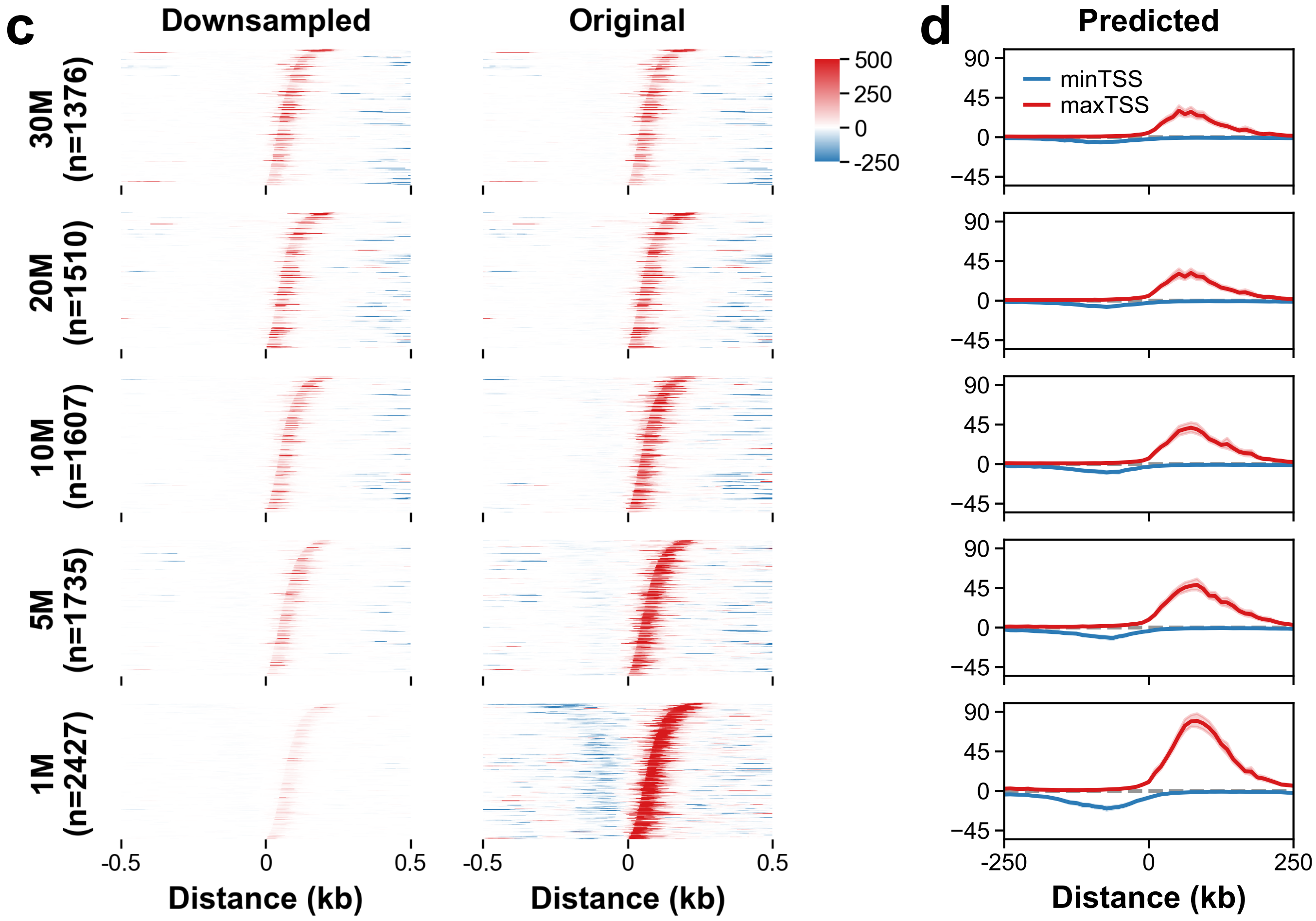

**Extended Data Fig. 3 | The effects of sequencing depth on binary classification of transcription directionality.**

**(a)** Observed PRO-cap signals of unidirectional distal elements at the corresponding downsampled levels (left) and at the original sequencing depth (right). The heatmaps show signals at individual elements, sorted by distance from the prominent TSS to the center.

**(b)** PRO-cap signals (10-bp bins) of unidirectional distal elements predicted by the ProCapNet model.

**(c)** Same as (a) for proximal elements.

**(d)** Same as (b) for proximal elements.

#### Extended Data Figure 4 | TEs & distal TREs

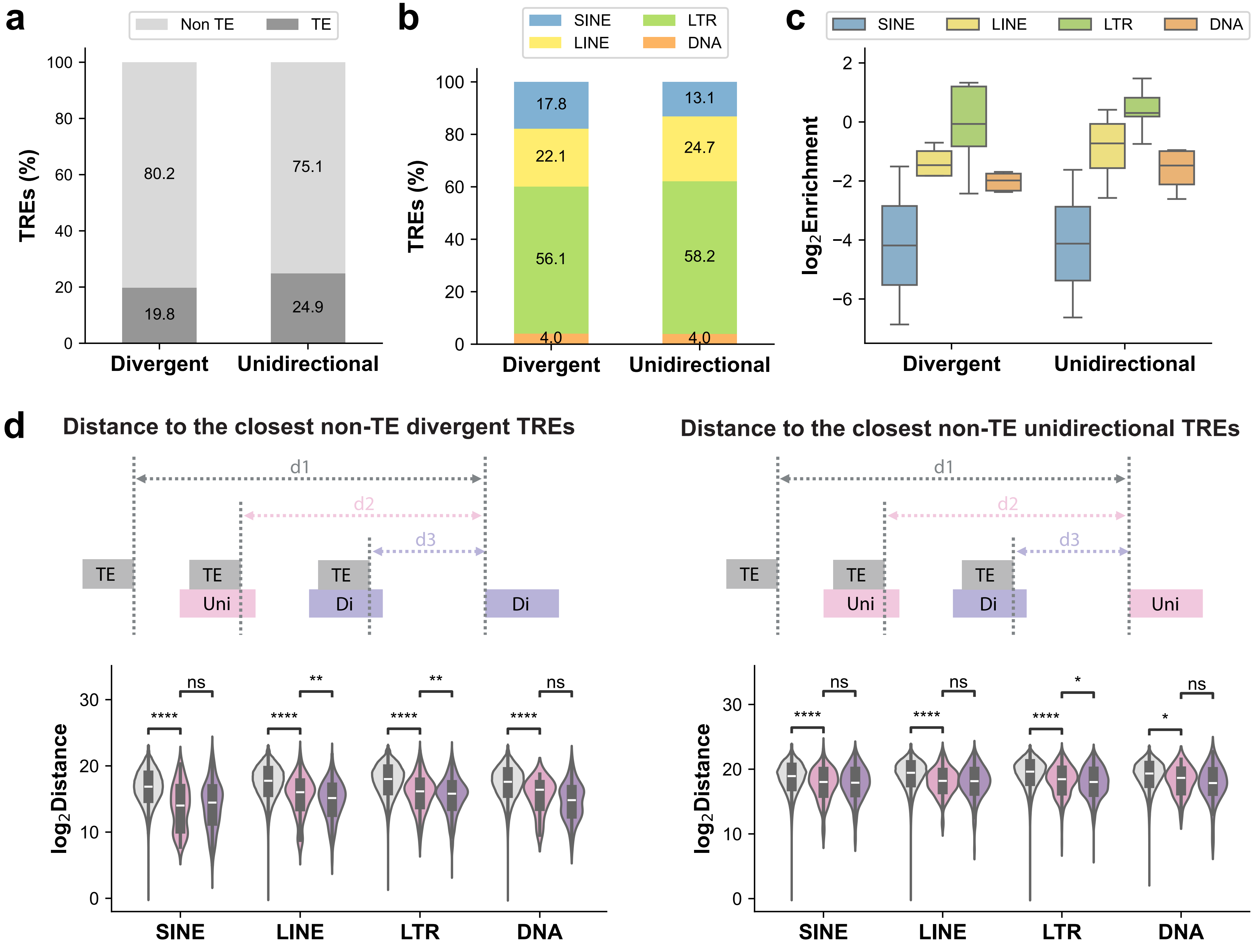

**Extended Data Fig. 4 | Association between TEs and distal elements.**

**(a)** Percentage distribution of TE- and non-TE-derived elements in divergent and unidirectional distal categories.

**(b)** Percentage distribution of each TE class (SINE, LINE, LTR, and DNA) found in TE-derived divergent and unidirectional distal elements.

**(c)** Enrichment of TE families grouped by TE class in divergent and unidirectional distal elements.

**(d)** Distance ($log_2$, bp) of TEs to the closest non-TE-derived divergent (left) and unidirectional (right) distal elements. $d_1$: the distance between TEs that do not overlap with TREs and the closest non-TE-derived distal elements. $d_2$: the distance between TEs that overlap with unidirectional distal elements and the closest non-TE-derived distal elements. $d_3$: the distance between TEs that overlap with divergent distal elements and the closest non-TE-derived distal elements.

#### Extended Data Figure 5 | Genomic distribution of distal elements

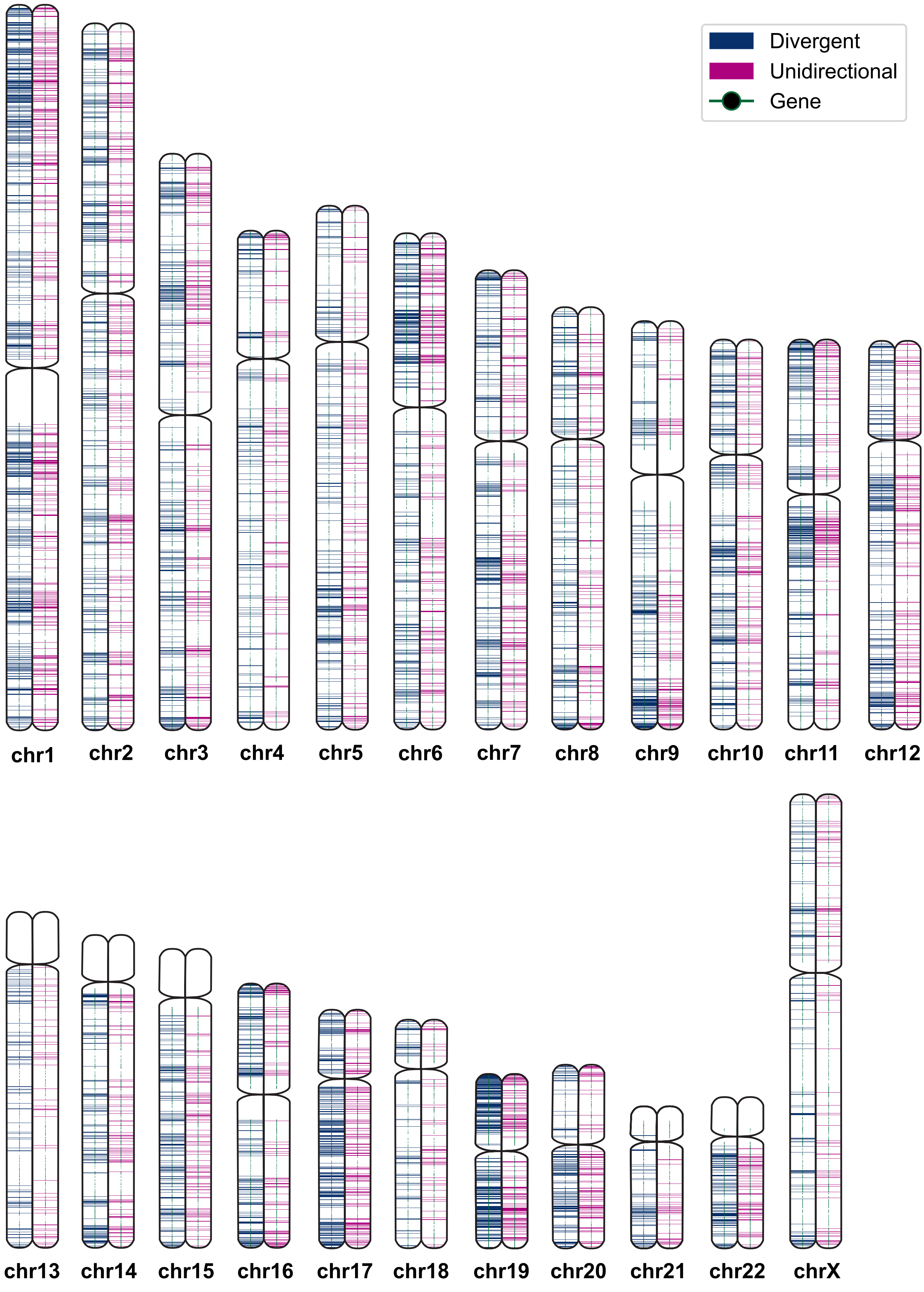

**Extended Data Fig. 5 | Genomic distribution of divergent and unidirectional distal elements.**

Distribution of divergent (blue) and unidirectional (pink) distal elements in 50-kb bins along the genome, with genes marked by green dots.

**Extended Data Figure 6 | Functional features and evolutionary metrics of distal elements**

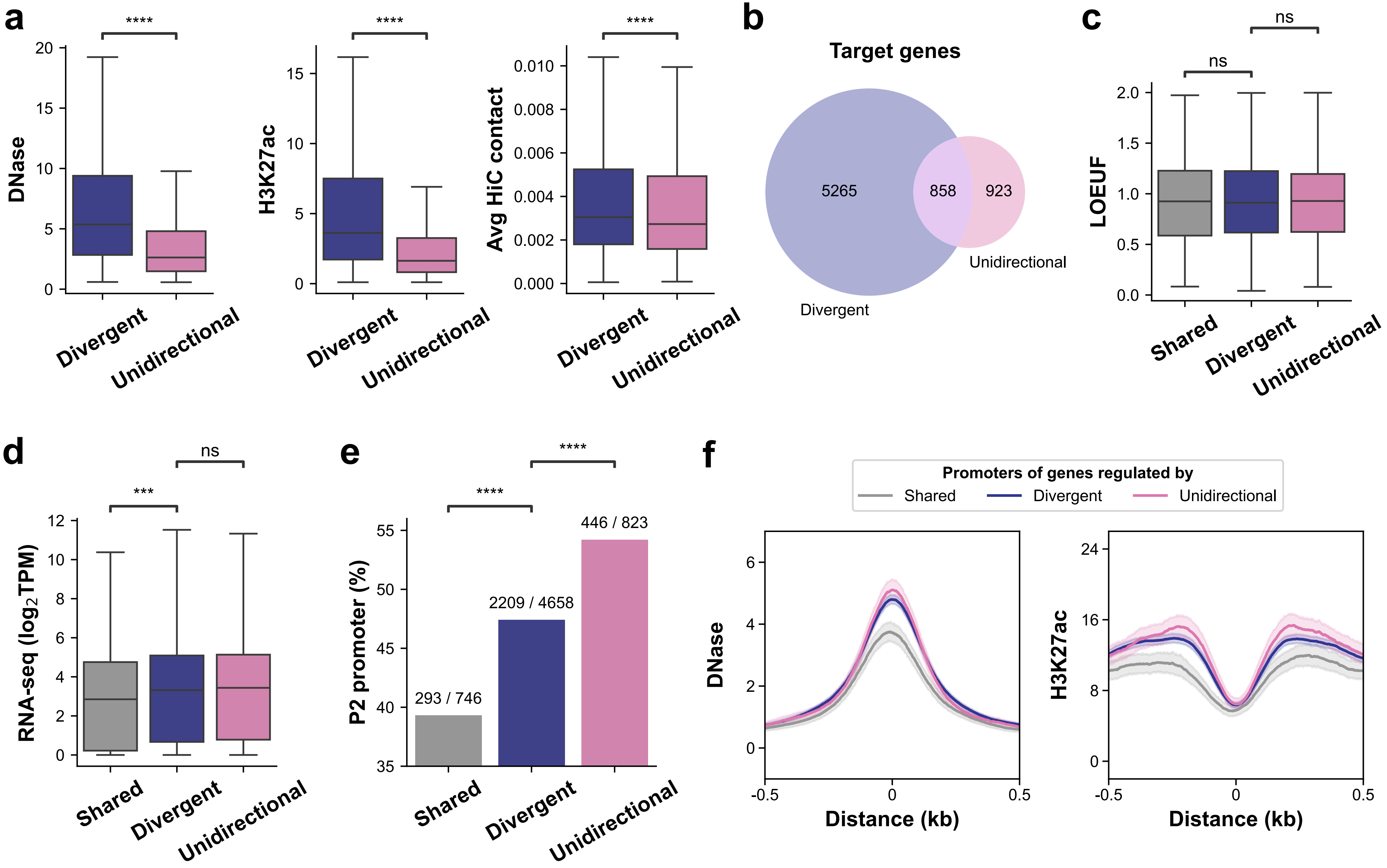

**Extended Data Fig. 6 | Functional features and evolutionary metrics of divergent and unidirectional distal elements.**

**(a)** Barplots of normalized DNase-seq and H3K27ac ChIP-seq values (with the geometric mean of these values serving as the activity component in the ABC score) and power-law scaled KR normalized Hi-C data (used as the contact component in the ABC score), averaged across all tested promoter-enhancer pairs for a given distal element.

**(b)** Venn diagram showing the number of predicted target genes unique to divergent elements, unique to unidirectional elements, or shared between both types.

**(c-f)** The distribution of LOEUF scores (c), gene expression level measured by RNA-seq (d), proportion of P2 promoters (e), and promoter activity based on DNase-seq and H3K27ac ChIP-seq (f) across target genes in each category. Distances in (f) are represented as ±0.5 kb from the gene TSS.

**Extended Data Figure 7 | CTCF binding blocks initiation on the minimum TSS side of distal elements**

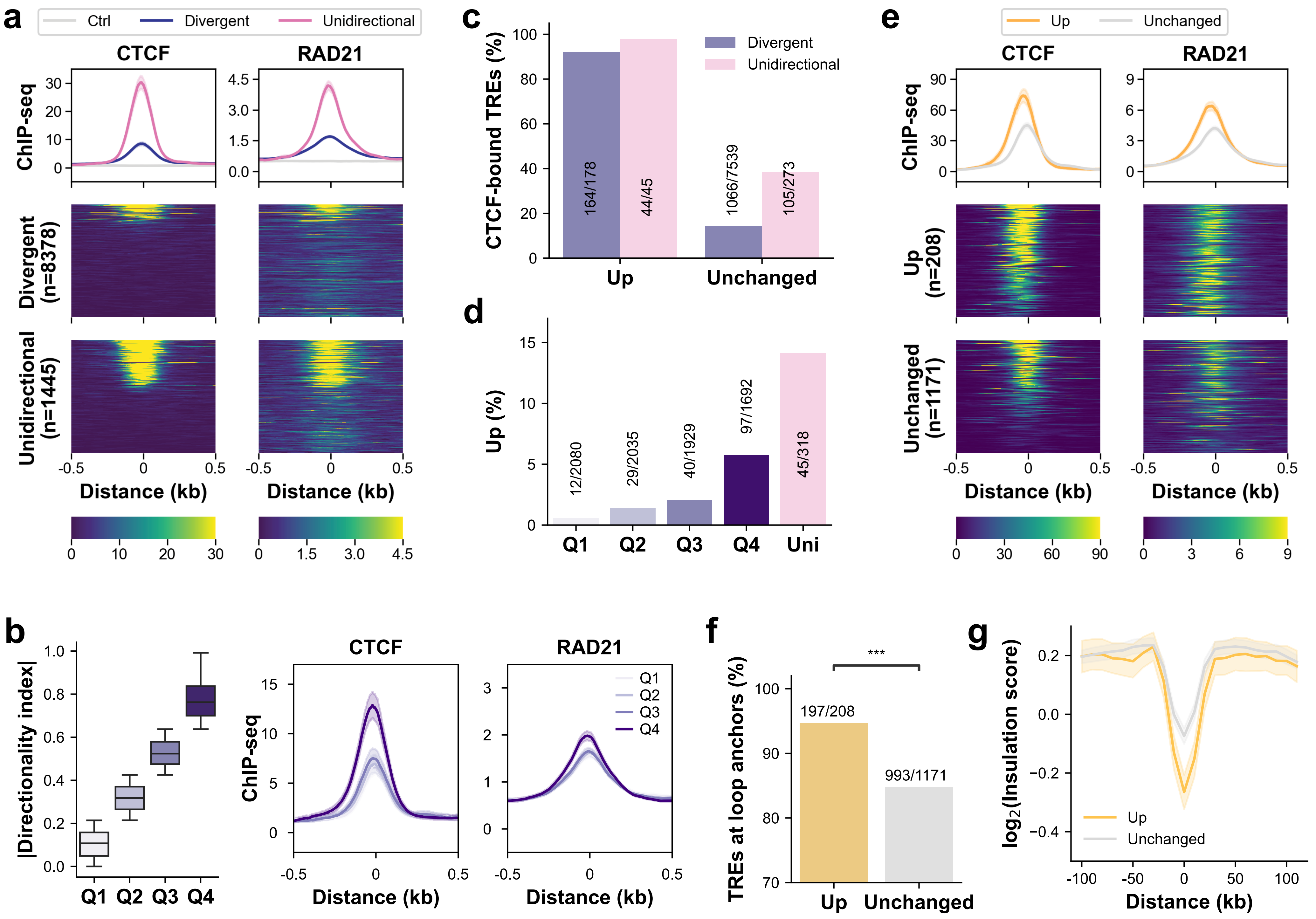

**Extended Data Fig. 7 | CTCF binding blocks initiation on the minimum TSS side of distal elements.**

**(a)** Metaplots and heatmaps of CTCF and RAD21 ChIP-seq signals at divergent and unidirectional distal elements. The elements in the heatmaps are sorted by CTCF ChIP-seq signals. Distances are represented as ±0.5 kb from the center.

**(b) Left:** Divergent distal elements are divided into four categories based on their absolute directionality index, with Q1 representing the most balanced elements and Q4 representing the most skewed. **Right:** Metaplots of CTCF and RAD21 ChIP-seq signals across different directionality categories.

**(c)** Percentage of elements with CTCF binding in divergent and unidirectional distal categories with or without transcription changes on the minimum TSS side.

**(d)** Percentage of elements with transcription upregulation after CTCF loss in unidirectional and divergent distal categories with varying directionality.

**(e)** Metaplots and heatmaps of CTCF and RAD21 ChIP-seq signals at CTCF-bound distal elements in transcriptionally upregulated and unchanged categories. The elements in the heatmaps are sorted by CTCF ChIP-seq signals. Distances are represented as ±0.5 kb from the center.

**(f)** Percentage of CTCF-bound distal elements located at chromatin loop anchors based on CTCF ChIA-PET in the upregulated and unchanged transcription groups.

**(g)** Metaplot of insulation scores across CTCF-bound distal elements, centered on CTCF motifs, in upregulated and unchanged groups.

#### Extended Data Figure 8 | CTCF's dual function is associated with directionality

**a**

| minTSS \ maxTSS | Up | Unchanged | Down |
|---|---|---|---|
| **Up** | 13 (5.8%) | 190 (85.2%) | 20 (9.0%) |
| **Unchanged** | 79 (1.0%) | 7548 (96.6%) | 184 (2.4%) |

**b**

| maxTSS \ minTSS | Up | Unchanged | Down |
|---|---|---|---|
| **Down** | 20 (9.3%) | 184 (85.6%) | 11 (5.1%) |
| **Unchanged** | 190 (2.5%) | 7548 (97.4%) | 8 (0.1%) |

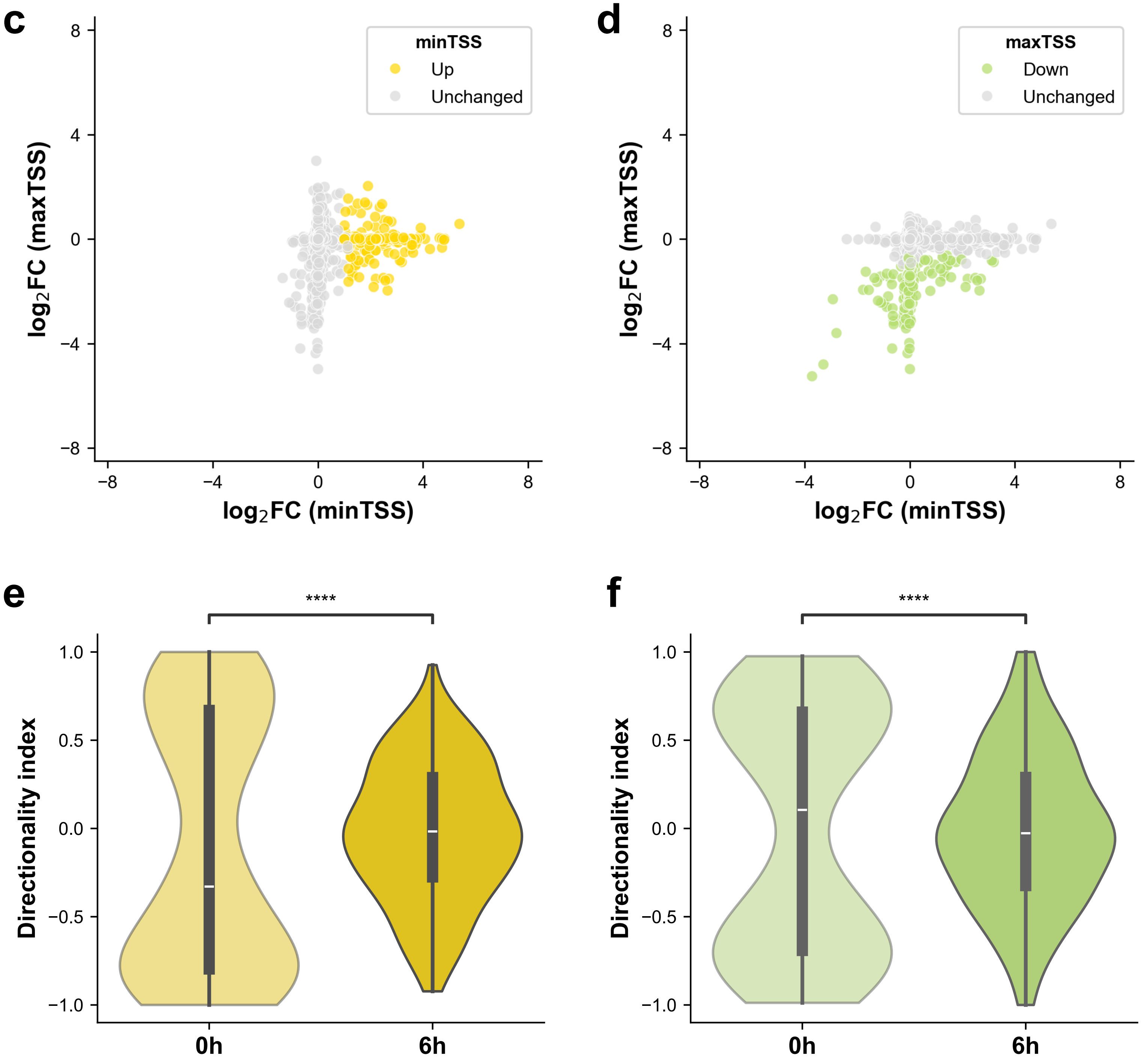

**Extended Data Fig. 8 | CTCF's dual function is associated with directionality.**

**(a)** Proportion of distal elements, categorized by transcription changes at the minimum TSS side, displaying upregulated, unchanged, and downregulated transcription patterns on the other side.

**(b)** Same as (a), but categorized by transcription changes at the maximum TSS side.

**(c)** Scatterplot comparing shrunken log fold changes in transcription level at the minimum and maximum TSS sides of the same distal elements. Data points are colored based on the changes at the minimum TSS side.

**(d)** Same as (c), but data points are colored based on the changes at the maximum TSS side.

**(e)** The distribution of directionality index before and after CTCF degradation among distal elements with upregulated transcription at the minimum TSS side.

**(f)** Same as (e), but for elements with downregulated transcription at the maximum TSS side.

**Extended Data Figure 9 | CTCF loss exposes optimal initiation sites**

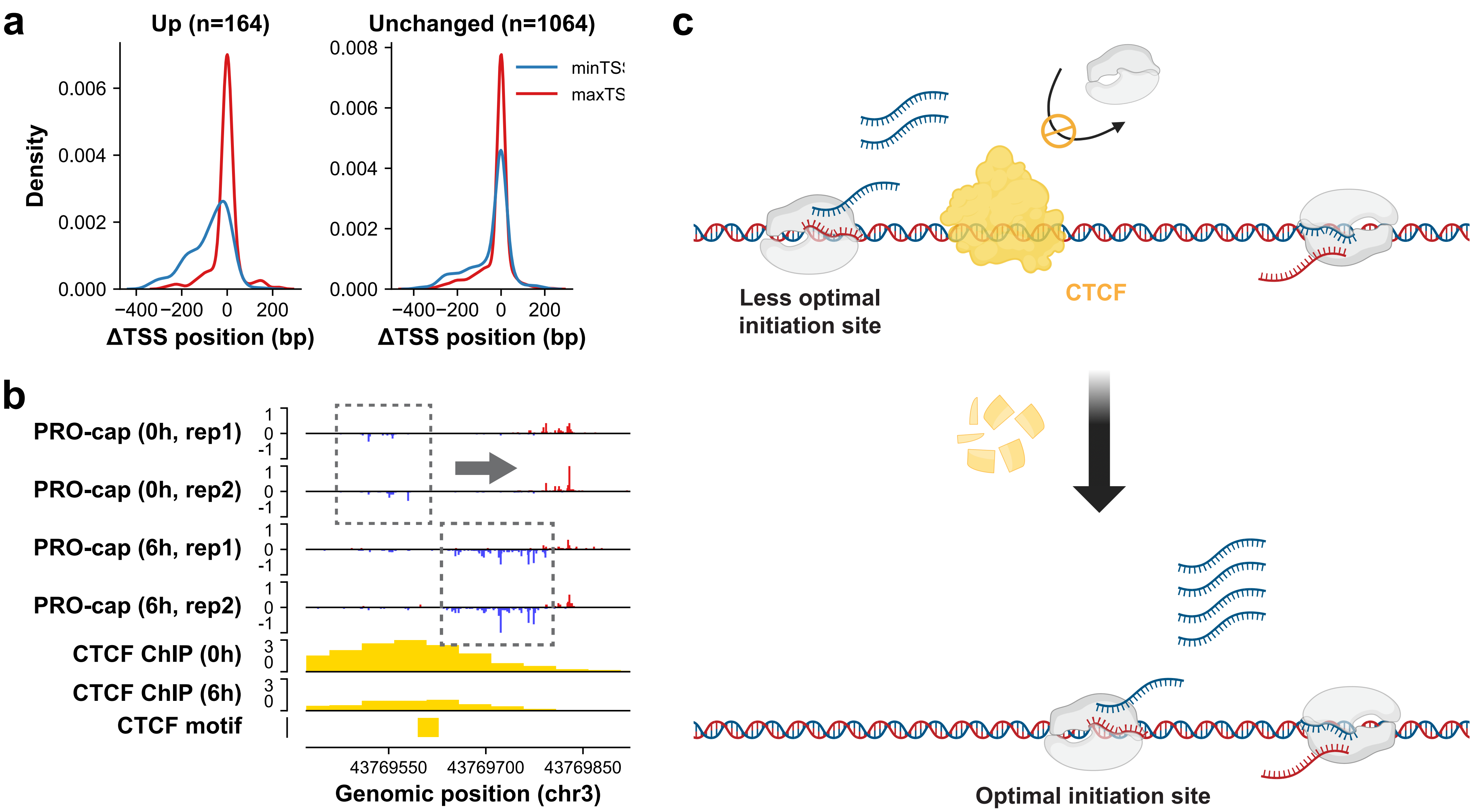

**Extended Data Fig. 9 | CTCF loss exposes optimal initiation sites.**

**(a)** Distribution of position changes for the minimum and maximum TSSs of the same elements after CTCF degradation in upregulated and unchanged transcription categories. Only CTCF-bound divergent distal elements are shown. Positive values indicate that TSSs have been repositioned downstream of their original locations, while negative values indicate an upstream shift.

**(b)** Genome browser shot of a representative distal element showing TSS repositioning toward the upstream region after CTCF degradation. The position of the CTCF motif is indicated and PRO-cap signal tracks from two replicates are displayed.

**(c)** Illustration of TSS repositioning after CTCF loss.

#### Extended Data Figure 10 | CTCF's dual function in proximal elements

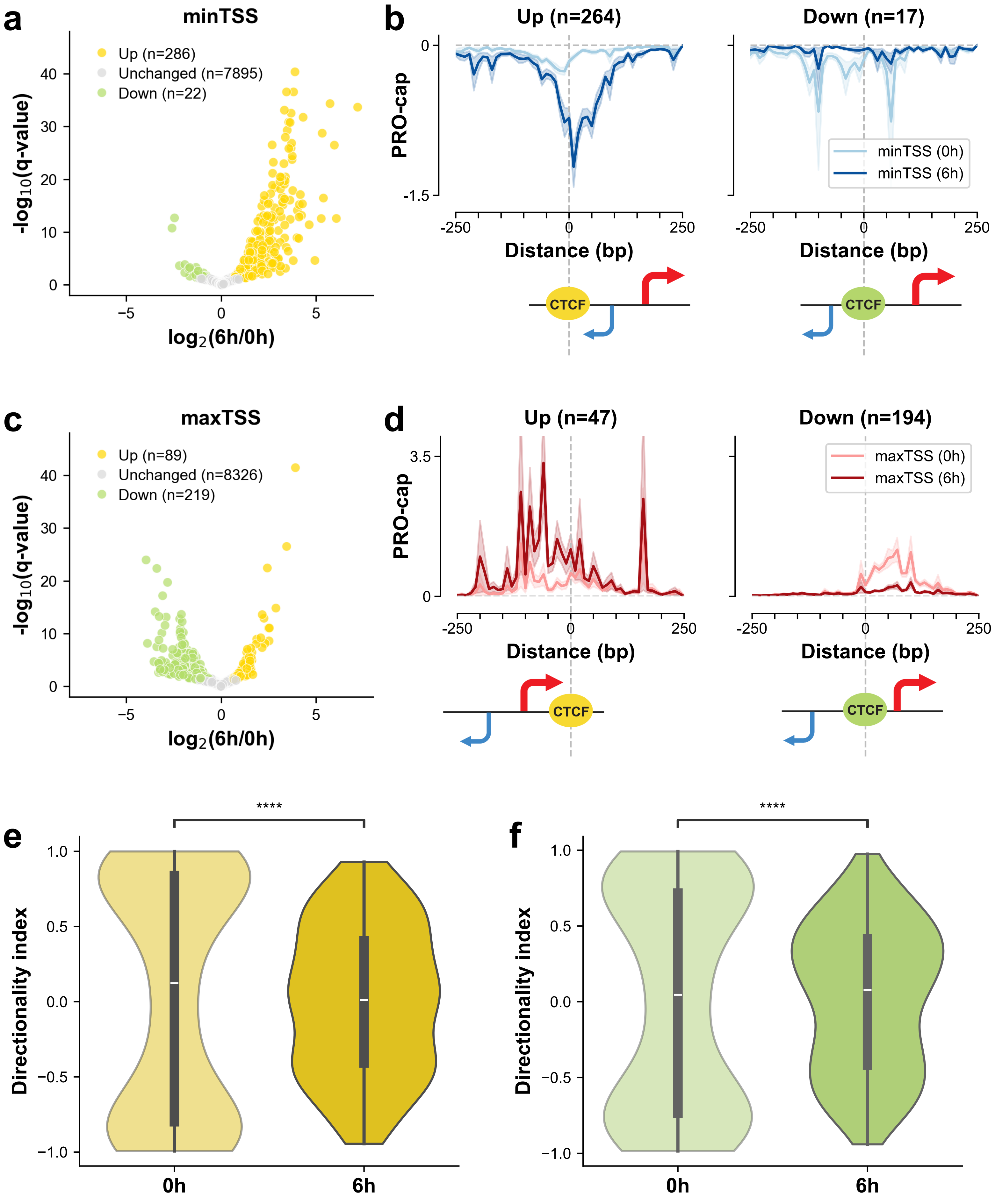

**Extended Data Fig. 10 | CTCF's dual function in proximal elements.**

**(a)** Volcano plot of PRO-cap signals on the minimum TSS side of proximal elements before and after CTCF degradation. Differentially expressed proximal elements are highlighted in yellow for upregulation and green for downregulation in transcription.

**(b)** Metaplots of PRO-cap signals (RPM normalized, 10-bp bins) on the minimum TSS side of proximal elements in the upregulated (left) and downregulated (right) transcription groups, centered on CTCF motifs. A schematic of the CTCF motif position relative to transcription initiation sites is shown below each panel.

**(c)** Same as (a), but for the maximum TSS side of proximal elements.

**(d)** Same as (b), but for the maximum TSS side of proximal elements.

**(e)** The distribution of directionality index before and after CTCF degradation among proximal elements with upregulated transcription at the minimum TSS side.

**(f)** Same as (e), but for elements with downregulated transcription at the maximum TSS side.

### Extended Data Figure 11 | ProCapNet predicts CTCF's dual functions

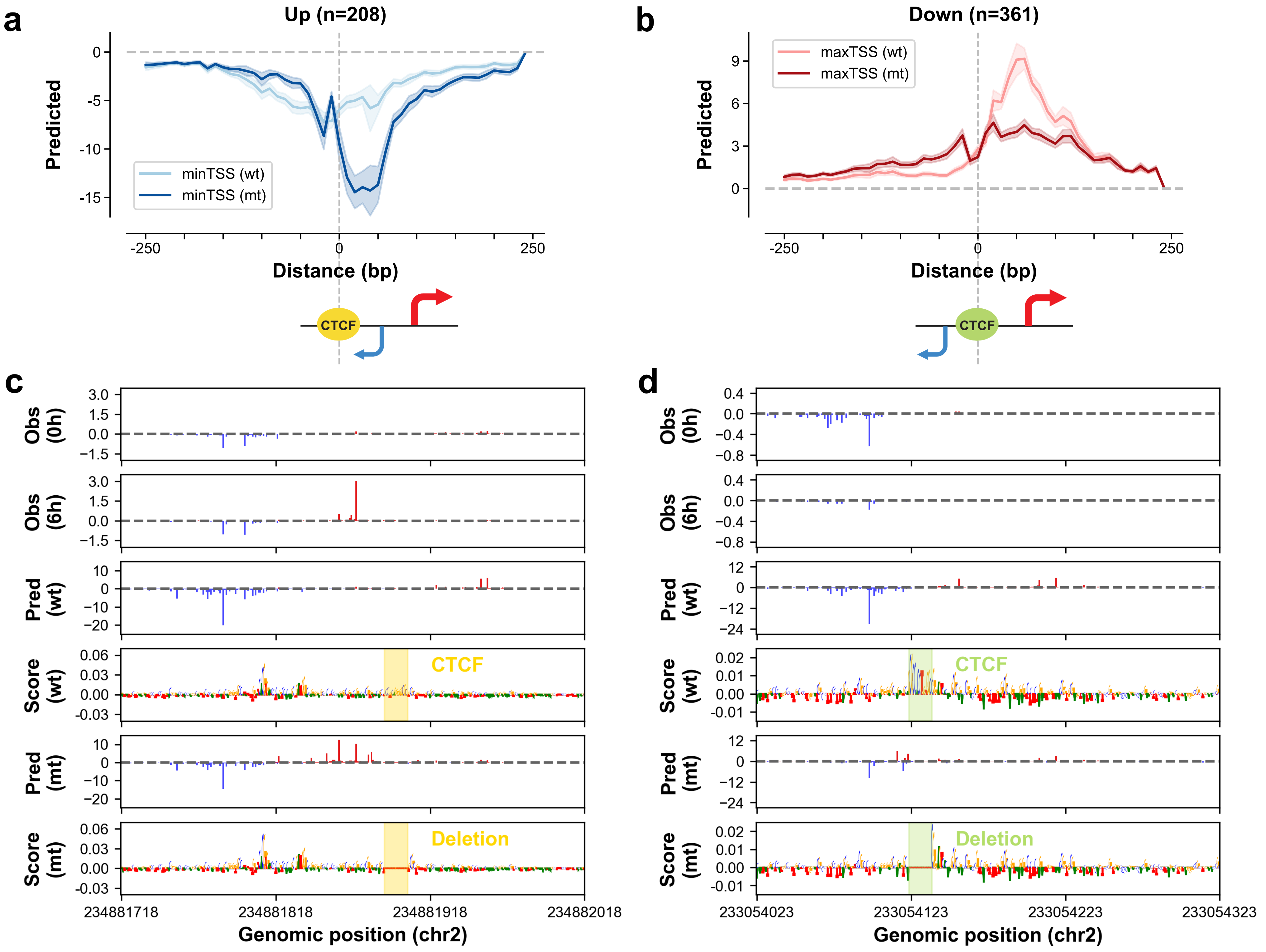

**Extended Data Fig. 11 | ProCapNet predicts CTCF's dual functions.**

**(a-b)** Predicted PRO-cap signals (10-bp bins) for distal elements with increased (a, same element set as Fig.4d-left) or decreased (b, same element set as Fig.4f-right) transcription levels in degron experiments, both before and after *in silico* deletion of CTCF motifs.

**(c-d)** Two representative loci where *in silico* deletion of CTCF motifs resulted in either transcription upregulation (c) or downregulation (d). The first two tracks display observed PRO-cap signals (RPM normalized) before and after CTCF degradation in degron experiments. The following four tracks present predicted PRO-cap signals and their corresponding contribution scores from the count task based on original and mutant sequences. CTCF motifs are highlighted in shaded areas.

Extended Data Figure 12 | Motifs with orientation- and position-dependent effects on transcription initiation

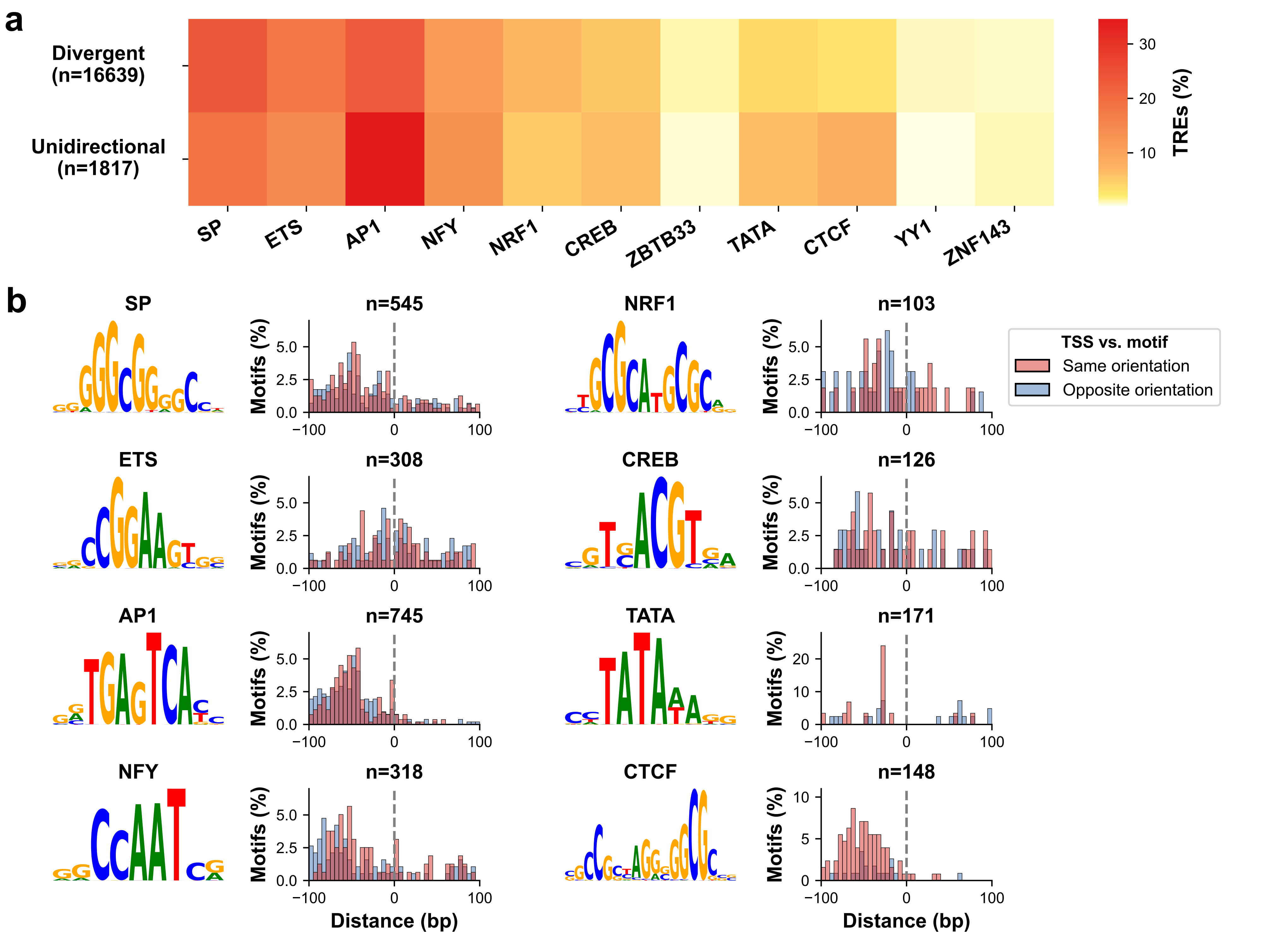

**Extended Data Fig. 12 | Motifs with position- and/or orientation-dependent effects on transcription initiation.**

**(a)** Heatmap showing the percentage of divergent and unidirectional elements with a given motif type that contribute to their transcription levels.

**(b)** The sequence logos represent the contribution weight matrix of each motif. The histograms show the distribution of motif positions relative to prominent TSSs of unidirectional elements, categorized by whether the TSS and motif align in the same or opposite orientation.
